# Tracking the intensity of the mechanism to produce antigenic diversity by subtelomeric ectopic recombination across the phylogeny of *Plasmodium* parasites

**DOI:** 10.1101/2023.02.27.530335

**Authors:** Carolina Martínez, Heiber Cárdenas, Mario A. Cerón-Romero

## Abstract

The generation of antigenic diversity, key for parasite virulence, has been investigated in the genus *Plasmodium*, mainly in *Plasmodium falciparum*. Cytogenetic and molecular studies have revealed that its subtelomeres are rich in antigenic gene families and undergo ectopic recombination. As a result, these families are highly variable and even species-specific. More recent analyses focused on the phylogenetic mapping of *P. falciparum* chromosomes with the bioinformatic tool PhyloChromoMap, showed that ectopic recombination of subtelomeres extends to all chromosomes. Although antigenic gene families have been described in subtelomeres of other *Plasmodium* species, the intensity of this mechanism in these species is still unclear. In this study, we investigated to what extent ectopic recombination of subtelomeres drives the generation of antigenic diversity in 19 *Plasmodium* species. To achieve this, we analyzed the profile of gene conservation in maps of all their chromosomes with PhyloChromoMap. Our results suggest that ectopic recombination of subtelomeres is more critical for the diversification of *pir* or *rif*/*stevor* genes than other antigenic gene families. Furthermore, its intensity varies among subgenera and was likely acquired and lost multiple times in the phylogeny of *Plasmodium*.

## 2 Introduction

The genus *Plasmodium* belongs to the clade Apicomplexa and includes more than 200 species of protozoan hemoparasites that use dipterans as vectors to infect a great diversity of vertebrate hosts. Phylogenetic analyses of these species show conflicts between their taxonomy and their phylogeny, as well as between their phylogeny and their host phylogeny (Galen, *et al*., 2018; Bo□hme, *et al*., 2018; Rich & Xu, 2011). For instance, the two most popular species affecting humans, *P. falciparum* and *P*.*vivax*, belong to two different subgenera. While *P. falciparum* belongs to *Laverania*, a subgenus that includes parasites of primates such as humans, gorillas, chimpanzees, and bonobos; *P. vivax* belongs to the subgenus *Plasmodium*, which includes parasites of primates not included in *Laverania* (Sharp, *et al*., 2020). In addition to *Plasmodium* and *Laverania*, the next most studied genera are *Haemamoeba*, which infects birds; and *Vinckeia*, which infects rodents (Pacheco, *et al*., 2011; Perkins, 2014; Sharp, *et al*., 2020). Although the phylogenetic order of these four subgenera has been a matter of debate, the most widely accepted proposal suggests that *Haemamoeba* diverges first, followed by *Laverania*, and finally, by the sister clades *Vinckeia* and *Plasmodium* (Borner, *et al*., 2016; Galen, *et al*., 2018; Escalante, *et al*., 2022).

The discordance between the phylogenies of *Plasmodium* species and those of their hosts suggests that these parasites have highly dynamic genomes and that their infection mechanisms have allowed them to frequently change and diversify their hosts (Galen, *et al*., 2018; Bo□hme, *et al*., 2018; Rich & Xu, 2011). Hence, comparative analyses of their genomes can reveal evolutionary patterns such as those related to their infection mechanisms. Although there are more than 30 annotated genomes and more than 200 described species of *Plasmodium*, most of the molecular and genomic studies have focused mostly on the species that infect humans, particularly *P. falciparum* and *P. vivax*. The former is the most virulent and deadly malaria agent, and the latter is the most widely distributed malaria agent worldwide (WHO, 2014).

Genomic characteristics of *P. falciparum* that have been compared with a few other species include the size of the genome, the number of chromosomes, and the structure of the subtelomeres. *P. falciparum* has a 23 Mb genome organized into 14 linear chromosomes ranging from 0.7 to 3.4 Mb (Hernández-Rivas, *et al*., 2013; Kemp, *et al*., 1987) and studies in other *Plasmodium* species, mainly those infecting mammals, show similar chromosomal organization and size (Carlton, *et al*., 1999; Pain, *et al*., 2008; Carlton, *et al*., 2008). Moreover, subtelomeres of *P. falciparum* are significantly less conserved than the chromosomic internal regions, a feature that is strongly linked to its virulence (Hernandez-Rivas, *et al*., 2013; Reed, *et al*., 2021), which has also been documented in *P. vivax* and *P. knowlesi* (del Portillo, *et al*., 2001; Pain, *et al*., 2008). Indeed, the high sequence variation observed in the subtelomeres of *P. falciparum* lies in sequences encoding virulence factors, while the rest of the subtelomeres and telomeres are composed of repeats that tend to be conserved (Scherf, *et al*., 2001; Hernández-Rivas, *et al*., 2013). These more specific details about subtelomeric and telomeric structures have been less explored in other *Plasmodium* species.

The importance of the chromosome structure in promoting antigenic variation as proposed in *P. falciparum* and *P. vivax* (Figueiredo, *et al*., 2002; del Portillo, *et al*., 2001) is also common in other eukaryotic taxa such as excavate parasites. However, both the mechanisms and the chromosomal regions involved are very variable (Silva Pereira, *et al*., 2020; Arkhipova & Morrison, 2001). In *P. falciparum* as well as in *Trypanosoma brucei* and *Trypanosoma cruzi*, the central mechanism for generating antigenic diversity is ectopic recombination of subtelomeres (Freitas-Junior, *et al*., 2000; Barry, *et al*., 2003). There is cytogenetic evidence that in *P. falciparum*, this subtelomeric recombination is facilitated by the anchoring of chromosomes at the nuclear periphery that brings subtelomeres closer (Freitas-Junior, *et al*., 2000). In *P. falciparum*, ectopic recombination of subtelomeres to generate antigenic diversity occurs because subtelomeres from non-homologous chromosomes share repeated sequences (Barry, *et al*., 2003). This process facilitates gene conversion events that produce new gene variants (Freitas-Junior, *et al*. 2000), resulting in large antigenic gene families that in many cases are specie-specific (Kooij, *et al*., 2005; Frank, *et al*., 2008; Otto, *et al*., 2018).

The evolutionary history of this mechanism of subtelomeric recombination to generate antigenic diversity is still unknown, but describing the intensity of this mechanism in each species and for each antigenic gene family can offer important clues to reconstruct such history. In *P. falciparum*, at least three subtelomeric multigene families have been documented, *var, rif* and stevor (Su, *et al*., 1995; Cheng, *et al*., 1998), while the *vir* family has been described in *P. vivax* (Bowman, *et al*., 1999). A recent study implementing a new phylogenomic mapping method on *P. falciparum* chromosomes with the tool PhyloChromoMap estimated that this molecular mechanism is the main mechanism for generating diversity in the *rif* and *stevor* families, but not in *var* (Cerón-Romero, *et al*., 2018). Although it is suggested that this mechanism is important for the diversification of the *vir* family of *P. vivax*, its level of intensity has not yet been determined (del Portillo, *et al*., 2001; Carlton, *et al*., 2008). In fact, the presence and intensity of this mechanism in other *Plasmodium* species and their multigene families have not been determined either. Those gene families and species are the *sicavar* and *kir* genes in *P. knowlesi* (Al-Khedery, *et al*., 1999; Janssen, *et al*., 2004), *cir* in *P. chabaudi, bir* in *P. berghei*, and *yir* in *P. yoelii* (Janssen, *et al*., 2002). While the *vir, kir, cir, bir* and *yir* families have been widely recognized as members of the *pir* superfamily, their relationship to the *rif/stevor* families is a matter of controversy (Janssen, *et al*., 2004; Cunningham, *et al*., 2010; Harrison, *et al*., 2020). Sequence similarity and phylogeny data determined *rif/stevor* as members of *pir* (Janssen, *et al*., 2004), while protein structure data reject this proposal (Harrison, *et al*., 2020). Contrasting information about the evolution of these gene families, the phylogenies of the parasites, and levels of intensity of this mechanism to promote antigenic diversity can also offer some clarity to this controversy.

Although molecular and phylogenetic chromosome mapping analyses have shown the importance of ectopic recombination of subtelomeres for *P. falciparum* virulence, the level of incidence of this mechanism in other *Plasmodium* species is still uncertain. This study evaluates whether ectopic recombination of subtelomeres is the preferred mechanism for generating antigenic diversity in 19 *Plasmodium* species. For this purpose, this study makes inferences about the intensity of recombination looking for genomic signatures such as young subtelomeric regions with a high density of antigenic gene families. The tool PhyloChromoMap was used to build chromosome maps of gene conservation and the results were contrasted against a robust phylogeny of *Plasmodium* to assess possible events in the evolutionary history of this mechanism in *Plasmodium* parasites. This study concludes the importance of this complex mechanism, which seems to be acquired and lost multiple times in the history of *Plasmodium*, is clade-specific and more associated with the genes *pir* and *rif*/*stevor*.

## 3 Materials and Methods

The methodology of this study was divided into five steps. The first step was the phylogenomic reconstruction of *Plasmodium*, used as a frame of reference in subsequent analyses. The second step consisted of the construction of chromosomal maps of gene conservation. This step provided information on the presence of young subtelomeric and internal regions that could be candidates for recombinant regions. The third step was the analysis of antigenic gene distribution along the chromosomes. In this step, we sought to determine whether the distribution of antigenic genes coincided with young internal and/or subtelomeric regions. For step four, the results from steps two and three were used to establish criteria to classify each *Plasmodium* species according to the intensity of subtelomeric ectopic recombination to promote antigenic variation (Supplementary Table S1). These criteria, together with the presence of antigenic sequences (≥3) in young regions allowed the determination of candidate regions to undergo ectopic recombination to generate antigenic diversity (hereinafter referred to as “CERAD regions”). In step five, the information obtained in step four was contrasted with the phylogeny from step one to find patterns of evolution of this mechanism of antigenic diversity production in the evolutionary history of *Plasmodium*.

### 3.1 Databases of whole genomes for phylogenetic reconstruction

A database of complete genomes was constructed for the phylogenetic reconstruction of the genus *Plasmodium* (Supplementary material). The sequences were obtained from PlasmoDB (http://PlasmoDB.org) and GenBank (https://www.ncbi.nlm.nih.gov/genbank/). This database is composed of 40 species, of which 35 are *Plasmodium* and one is *Hepatocystis*, the parasite of the red colobus monkey *Piliocolobus* tephrosceles (Aunin, *et al*., 2020). *Plasmodium* species are distributed into four distinct subgenera (Escalante, *et al*., 2022; Perkins, 2014) as follows: *Laverania (*12), *Haemamoeba* (2), *Plasmodium* (14) and *Vinckeia* (7). On the other hand, the remaining four species (i.e., *Babesia bovis, Babesia bigemina, Theileria equi* and *Theileria annulata*) form the outgroup.

These species were also used as an outgroup in previous phylogenetic studies about *Plasmodium* (Salomaki, *et al*., 2021; Perkins & Schall, 2002; Borner, *et al*., 2016).

### 3.2 Reconstruction of the phylogeny of *Plasmodium*

Six different phylogenetic approaches were used to reconstruct the species tree used as a phylogenetic framework to compare conservation profiles among *Plasmodium* species. The first approach involved using OrthoFinder v2.5.4, which uses protein sequences as input to identify gene families, infer orthologs, and construct a species tree based on those orthologs (Emms & Kelly, 2015). The gene families obtained in the intermediate steps of OrthoFinder were used as input files for the other phylogenetic approaches of species tree reconstruction.

The remaining five approaches for species tree reconstruction consist of four summary gene tree analyses (Zhang, *et al*., 2020; Willson, *et al*., 2022; Morel, *et al*., 2022), three of which include multiple-copy gene families and one includes only single-copy genes; and a supermatrix analysis by alignment concatenation (de Queiroz & Gatesy, 2007). For all of these approaches, gene families present in all the taxa were chosen to be aligned with the *einsi* method (--ep 0 --genafpair -- maxiterate 1000) of MAFFT v7.505 (Katoh, *et al*., 2005). Then, PAL2NAL v14.0 was used to obtain the codons alignments (Suyama, *et al*., 2006). With the results from PAL2NAL, the phylogeny of each gene family was reconstructed using IQ-TREE v1.6.9 (Nguyen, *et al*., 2015), with parameters - B 1000 -alrt 1000 to obtain branch support (i.e., UFBoot and SH-aLRT) in all trees (Minh, *et al*., 2013; Guindon, *et al*., 2010). These phylogenetic trees were used to generate a species tree with three tools that use a set of multi-copy gene families as input file: ASTRAL-Pro v1.8.1.3 (Zhang, *et al*., 2020), ASTRAL v5.7.8-DISCO v1.3 (Willson, *et al*., 2022) and SpeciesRax v2.0.4 (Morel, *et al*., 2022). An additional species tree was reconstructed with ASTRAL v5.7.8 (Rabiee, *et al*., 2019) from single-copy gene trees. The alignments of these single-copy gene families were concatenated using Mega X v10.2.6 (Stecher, *et al*., 2020), and the resulting supermatrix was used for the inference of another species tree with IQ-Tree (Nguyen, *et al*., 2015).

Finally, the quality of the phylogenetic reconstructions of the species tree was evaluated, and the six versions of these trees were compared to each other and against other previously published versions (e.g., Escalante, *et al*., 2022; Galen, *et al*., 2018; Borner, *et al*., 2016). For the quality assessment, in addition to the analysis of branch support in the species trees, the median node support values were also analyzed in the 823 gene family trees used as input for the species tree inference. Furthermore, a majority rule consensus tree of the six species trees generated was constructed using PAUP* v4.0a168 (Swofford, 2003; for the full trees see supplementary material).

### 3.3 Databases for chromosome mapping of gene conservation

A complete genome database was constructed with 134 species distributed throughout the SAR clade (Stramenopiles, Alveolata, Rhizaria) and Excavata (Discoba and Metamonada) (Supplementary Table S2, Supplementary material). To have a higher resolution in Alveolata (Apicomplexa, Ciliophora and Dinozoa), sampling was focused on this clade. The rest of the sampling was evenly distributed between Stramenopila and Rhizaria, and 26 Excavata species (15 Discoba and 11 Metamonada) were also included. The selection of these genomes aimed to maximize the phylogenetic diversity of the group and the host diversity based on previous literature (Galen, *et al*., 2018; Pacheco, *et al*., 2018). For each of the 134 species, we sought to obtain protein sequences and if possible, coding sequences. These sequences were collected from different databases such as PlasmoDB (http://PlasmoDB.org), ToxoDB (https://toxodb.org), GenBank (https://www.ncbi.nlm.nih.gov/genbank/), PiroplasmaDB (https://piroplasmadb.org), CryptoDB (https://cryptodb.org), FungiDB (https://fungidb.org) and TriTrypDB (https://tritrypdb.org).

The protein and coding sequences database includes 33 species of *Plasmodium* distributed in four different subgenera (Escalante, *et al*., 2022; Perkins, 2014): *Laverania (*12), *Haemamoeba* (2), *Plasmodium* (12) and *Vinckeia* (7). Of these 33 species, 19 were reference genomes that contain annotated chromosomes and were selected to analyze their gene conservation profile along their chromosomes. This analysis was done considering the classification of the 19 species into the four *Plasmodium* subgenera: *Laverania (*7), *Haemamoeba* (1), *Plasmodium* (7), and *Vinckeia* (4).

The protein sequences of the 134 species were organized into gene families using OrthoFinder v.2.5.4. To do this, instead of running a complete analysis for inferring orthologs, these sequences were analyzed using the “tree only” configuration (*-ot*) of OrthoFinder. With this configuration, OrthoFinder infers homology among sequences, builds a database of gene families (referred as orthogroups by OrthoFinder), and reconstructs the phylogeny of each gene family (Emms & Kelly, 2015). Subsequently, DISCO v1.3 (Willson, *et al*., 2022) was used to resolve multi-copy gene families, and the resulting (resolved) single-copy gene families were used for running PhyloChromoMap (Cerón-Romero, *et al*., 2018).

### 3.4 Construction of chromosome maps of gene conservation

For the construction of the chromosome maps of the 19 *Plasmodium* species, we used PhyloChromoMap (https://github.com/marioalbertocer/PhyloChromoMap_py) with the phylogenetic trees generated in the previous section as input. The gene family mapping file required for PhyloChromoMap was prepared with information from PlasmoDB. Finally, for the centromere mapping file (also required for PhyloChromoMap), a custom Python script was created with the sliding window method that locates centromeres as the largest chromosomal region with the highest AT content, a feature that has been reported in some *Plasmodium* species in previous studies (Gardner, *et al*., 2002; Hoeijmakers, *et al*., 2012). After analyzing and comparing candidate centromere regions, distribution plots of AT content on chromosomes, and previous records in PlasmoDB, chromosome 2 of *P. cynomolgi* and *P. coatneyi*, chromosome 12 of *P. relictum* and chromosome 2 and 6 of *P. vivax-like* were left without a defined centromere (see results).

### 3.5 Identification and analysis of young chromosomic regions

Subtelomeric and internal young regions were defined as distinctive portions of the phylogenomic chromosome maps with low gene conservation, which was determined using a custom Ruby script and visual inspection. The script obtains candidate young regions that include a maximum of one conserved gene (present in three or more major clades). Initially, a standard maximum size of 200 kb and a standard minimum size of 80 kb were determined for young regions in all species, based on what was observed in *P. falciparum*, where all chromosomes have subtelomeric young regions at both ends (Cerón-Romero, *et al*., 2018). However, after visual inspection of the chromosome maps, it was determined that the size of young regions could be outside this range, so the size of some young regions was modified. In addition, each young region was manually reviewed to evaluate the presence of species-specific or young genes.

### 3.6 Analysis of the distribution of antigenic genes

Since the production of antigenic diversity through ectopic recombination is expected to generate young chromosomic regions with a high density of antigenic genes, the predominant gene families in young regions were sampled and their function was determined. For this, a custom Python script was used to identify the ten most frequent gene families in the young regions of each of the 19 *Plasmodium* species. The script also compares the incidence of these ten gene families in young subtelomeric, young internal, and conserved regions. We verified if these genes were antigenic by reviewing the literature on their products (Supplementary material). Subsequently, the antigenic genes were located on the chromosome maps to analyze their physical distribution (Supplementary material).

## 4 Results

### 4.1 Phylogenetic reconstruction of *Plasmodium*

OrthoFinder detected 7.415 gene families of which 823 present all taxa and generated an alignment and a phylogenetic tree. Of these 823 gene families, 597 are single-copy and 226 are multi-copy.

*Laverania, Vinckeia* and *Plasmodium* have a mean ratio of 1,137, 1,136 and 1,128 sequences per gene family respectively, and *Haemamoeba* (*P. relictum*) has 1.08 sequences per gene family. The quality of the 823 phylogenetic trees generated with IQ-Tree was good, since 99% of the gene families had a median bootstrap value between 80 and 100.

Phylogenomic analysis of *Plasmodium* showed a highly consistent topology among the species tree reconstruction approaches (Fig. 1C-G) and suggests that the subgenus *Plasmodium* is a non-monophyletic group (Fig. 1A). Only the tree generated by OrthoFinder shows *Plasmodium* as a monophyletic group, but with low branch support (<0.50, Fig. 1B). In contrast, the remaining five trees showed *Plasmodium* as non-monophyletic and at the base of the tree, with high branch support (>0.80, Fig. 1C-G). This topology also has important implications for *Hepatocystis* and *Vinckeia*, which appear in the early bifurcations of the OrthoFinder tree (Fig. 1B), whereas in the other five phylogenetic trees (Fig. 1C-G) they shared a most recent common ancestor and form the sister clade of *Laverania-Haemamoeba*. Finally, although the monophyly of *Plasmodium* is not supported by these results, seven of its taxa (*P. coatneyi, P. inui, P. fragile, P. knowlesi*, P. *cynomolgi, P. vivax* and *P. vivax-like*) form a recurrent monophyletic clade in the species trees, except in the tree generated by the supermatrix method (Fig. 1G).

**Figure 1.**
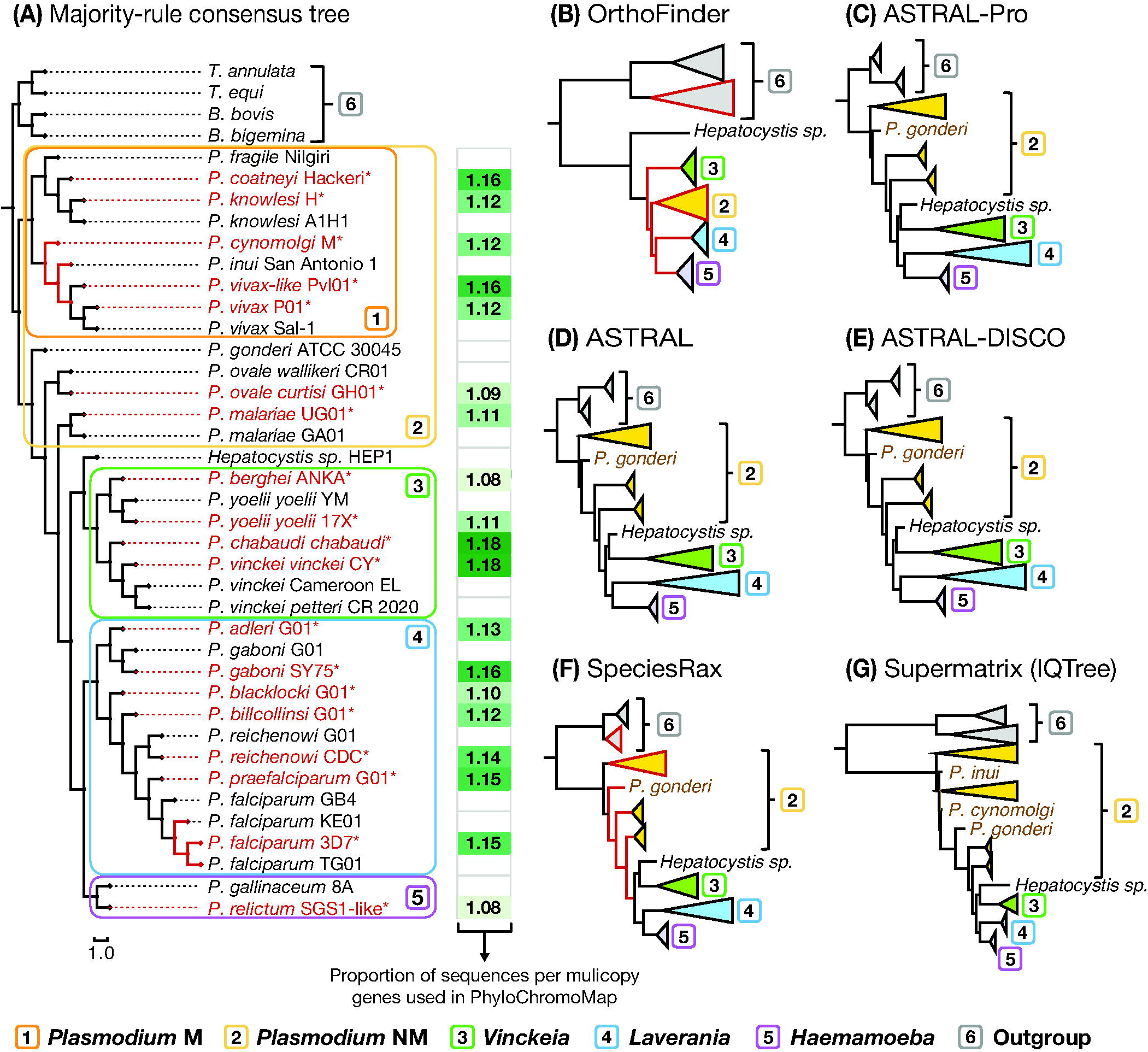
Phylogenetic reconstruction of the genus *Plasmodium*. **(A)** Majority-rule (50%) consensus tree generated with PAUP*, suggesting that the *Plasmodium* subgenus is not monophyletic. The proportion of sequences per gene family obtained with OrthoFinder, used in PhyloChromoMap, is shown next to each species. **(B)** OrthoFinder, **(C)** ASTRAL-Pro, **(D)** ASTRAL, **(E)** ASTRAL-DISCO, **(F)** SpeciesRax, (**G)** Supermatrix-IQTree. Red branches indicate low statistical support: Consensus (<80%), OrthoFinder (STAG consensus <70%), SpeciesRax (EQPIC<0,2). For the ASTRAL-Pro, ASTRAL, and ASTRAL-DISCO trees, all branches showed 100% support (LPP), except between *P. falciparum* strains 3D7 and TG01 (LPP=95-97%). All the branches in the supermatrix tree showed high support (SH-aLRT≥99%, UFBoot≥90%). Complete trees can be found in the supplementary material. *B. bigemina, B. bovis, T. annulata*, and *T. equi* were used as an outgroup to estimate the root of the trees. (*) = Species whose gene conservation profile was analyzed. *Plasmodium* NM = non-monophyletic (all species), *Plasmodium* M = monophyletic (monophyletic subgroup excluding *P. gonderi, P. ovale* and *P. malariae*).

### 4.2 Gene conservation profiles

OrthoFinder produced a database of 63.661 multi-copy gene trees, from which a database of 31.260 single-copy trees was produced using DISCO. Although about half of the tree database was discarded by DISCO (Supplementary Fig. S1), these trees did not meet the criteria of taxa inclusion of this tool (i.e., 50% of the trees were species-specific). Hence, the number of trees per species remained close to the original in each major clade: Apicomplexa 98%, Other alveolates 72%, Stramenopila 94%, Rhizaria 90%, Discoba 91%, and Metamonada 77% (Supplementary Fig. S1). Overall, between 25-50% of their phylogenetic trees were discarded for less than 13% of the species, and between 40-50% of the gene families were discarded for less than 5% of the species.

Phylogenetic chromosome maps showed that *Vinckeia* exhibits a distinctive gene conservation pattern, more consistent with the recombination of subtelomeres, which is significantly different from *Plasmodium* and *Laverania* (Fig. 2A-C). This pattern was characterized by young subtelomeres at almost all chromosome ends, and few young internal regions, which do not exceed 85 kb (Fig. 3).

**Figure 2.**
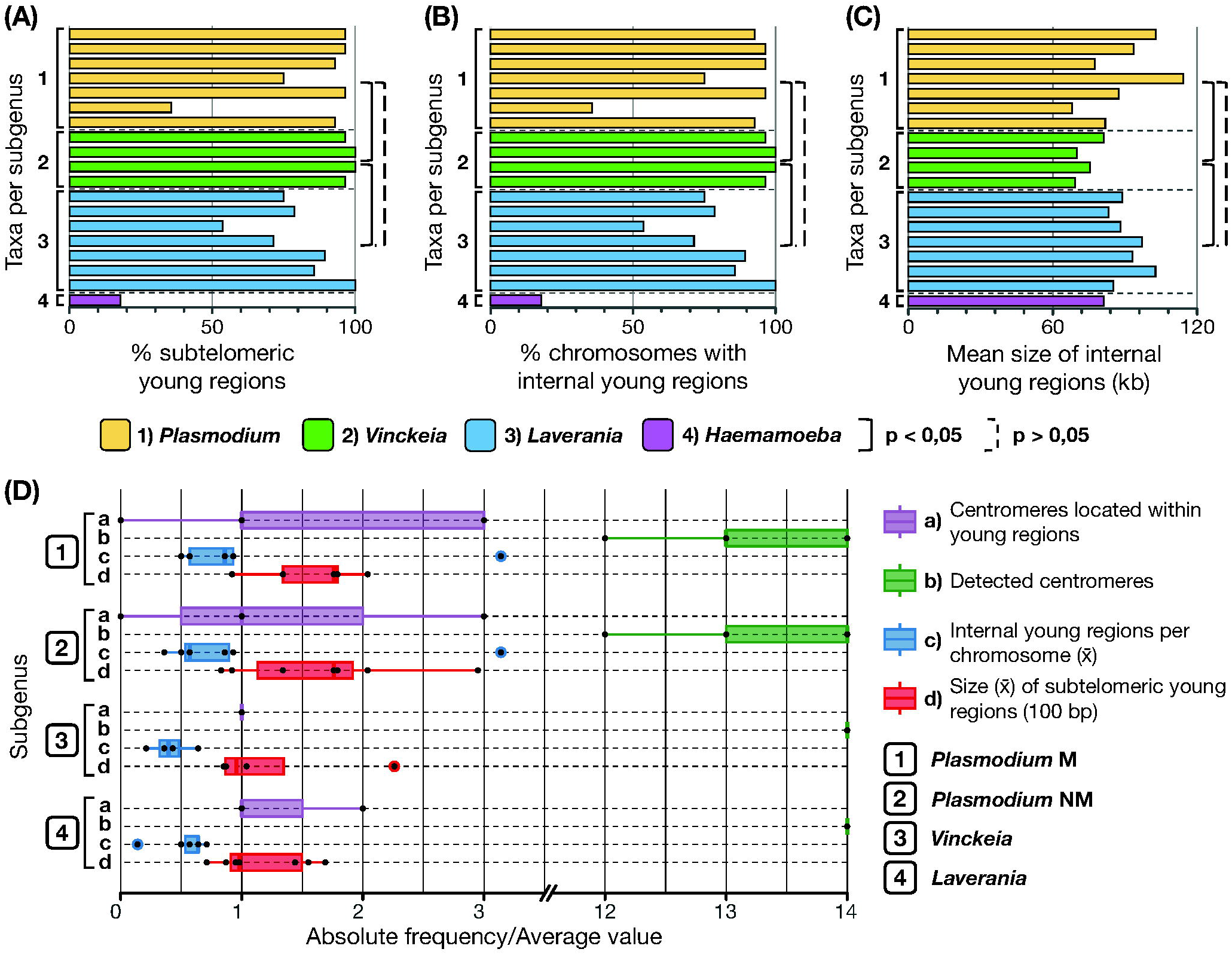
Characteristics of gene conservation profiles among subgenera of *Plasmodium*. **(A)** Percentage of young subtelomeres. *Vinckeia* shows a significantly higher percentage than *Laverania* (Wilcoxon-Mann-Whitney, W = 25, p = 0,0223) and *Plasmodium* (Wilcoxon-Mann-Whitney, W = 25, p = 0,0182). **(B)** Percentage of chromosomes with young internal regions. *Vinckeia* shows significantly lower values than *Laverania* (Wilcoxon-Mann-Whitney, W = 4,5, p = 0,0401) and *Plasmodium* (T-Student, t = -2,6039, p = 0,0143). **(C)** Average size of young internal regions (kb). *Vinckeia* shows significantly smaller values than *Laverania* (T-Student, t = -4,1503, p = 0,0012) and *Plasmodium* (T-Student, t = -1,874, p = 0,0469). **(D)** Characteristics with higher variation in *Plasmodium* than in *Vinckeia* and *Laverania*, where even the monophyletic clade of *Plasmodium* shows a higher variation. *Plasmodium* NM = non-monophyletic (all species), *Plasmodium* M = monophyletic (monophyletic subgroup).

**Figure 3.**
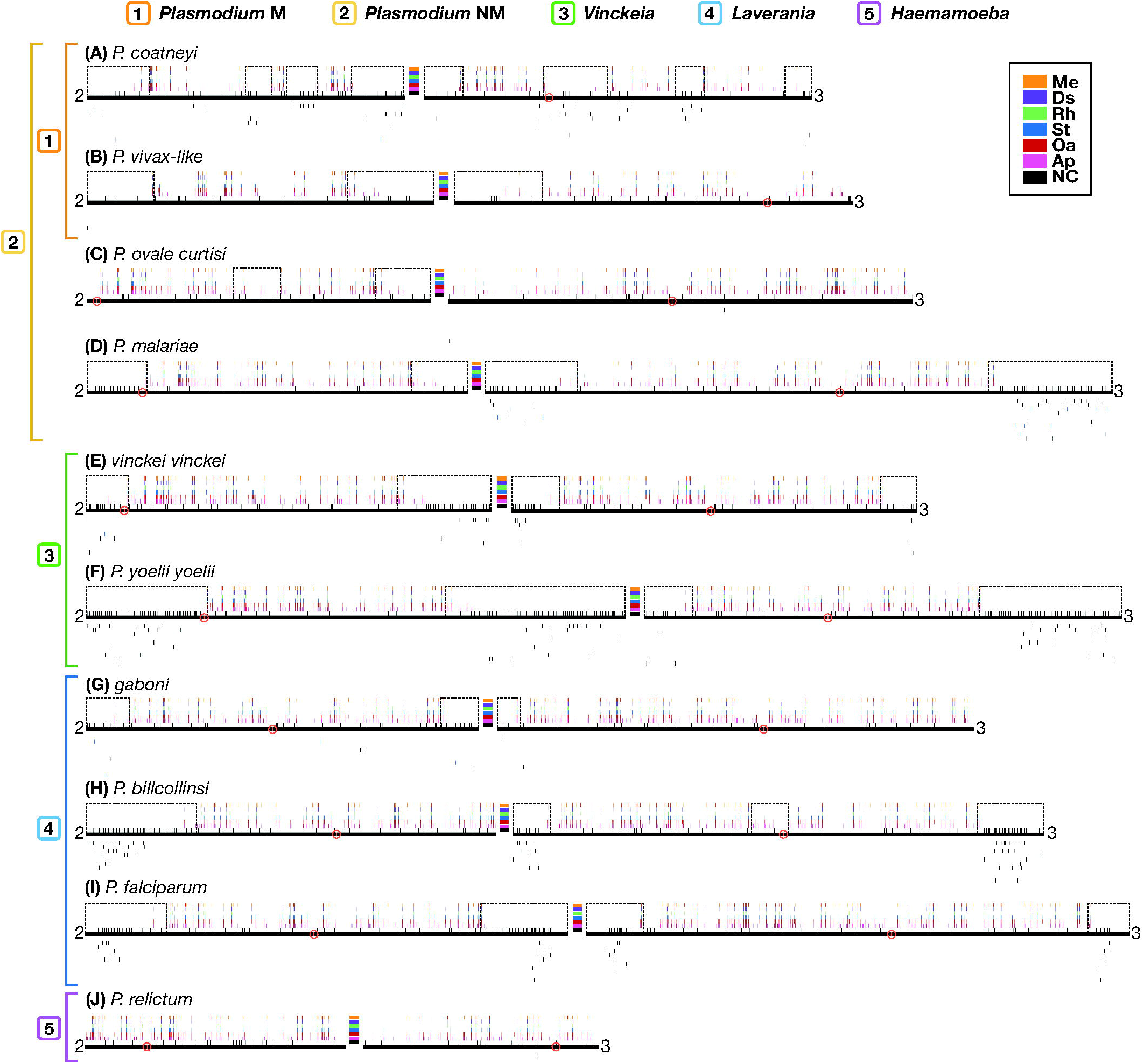
Examples of chromosome maps showing the gene conservation profile, presence of young regions, and distribution of antigenic genes in ten *Plasmodium* species distributed in *Plasmodium, Vinckeia, Laverania*, and *Haemamoeba*. Black lines represent chromosomes and bars above reflect levels of conservation, with dashed boxes around “young” regions. Detected centromeres are indicated by a red circle. Above the black line, the first row (NC) indicates genes whose phylogenetic trees do not meet the criteria of having more than ten taxa. The remaining rows (bottom to top) are heatmaps reflecting the proportion of lineages of Apicomplexa (Ap), Other Alveolates (Oa), Stramenopila (St), Rhizaria (Rh), Discoba (Ds), and Metamonada (Me) that contain the indicated gene. Lines below the chromosomes show the location of sequences belonging to antigenic gene families (black) or candidate antigenic gene families (blue), one per row, found in each species. *Plasmodium* NM = non-monophyletic (all species), *Plasmodium* M = monophyletic (monophyletic subgroup).

Accordingly, the proportion of young subtelomeres in *Vinckeia* is significantly higher than in *Laverania* (Wilcoxon-Mann-Whitney, W = 25, p = 0.0223) and *Plasmodium* (Wilcoxon-Mann-Whitney, W = 25, p = 0.0182). Likewise, the proportion of chromosomes with young internal regions and the average size of these regions was significantly lower in *Vinckeia* than in *Laverania* (Wilcoxon-Mann-Whitney, W = 4.5, p = 0.0401; T-Student, t = -4.1503, p = 0.0012, respectively), and *Plasmodium* (T-Student, t = -2.6039, p = 0.0143; T-Student, t = -1.874, p = 0.0469, respectively). Unlike *Vinckeia*, the subgenus *Plasmodium* showed high chromosomal structural variation among its species, even in those that are part of its monophyletic clade, and *Laverania* showed an intermediate pattern of variation compared to what was observed in *Vinckeia* and *Plasmodium* (Fig. 2D). On the other hand, *Haemamoeba* (*P. relictum*) exhibits less than 20% of chromosome ends as subtelomeres and less than 20% of chromosomes with young internal regions whose size is less than 80 kb (Fig. 2A-C).

### 4.3 Distribution of antigenic gene families

The search for the ten predominant gene families in young regions per species resulted in a total of 133 gene families (Supplementary material). In this process, *P. relictum* was the only species with less than ten families found (Fig. 4A). Out of the total number of gene families obtained, 11 were excluded from the analysis because it was not possible to verify whether their sequences were antigenic. Also, 14 gene families were classified as candidate antigenic genes since their sequences seem to play a key role in the virulence of these parasites, but their antigenic role could not be confirmed. Following this classification, most of the gene families by species (>80%; for the list of genes see Supplementary material), were suitable for the analysis of distribution on the chromosome maps, except in *Haemamoeba* (*P. relictum*) where 57% of its gene families were discarded (Fig. 4A).

**Figure 4.**
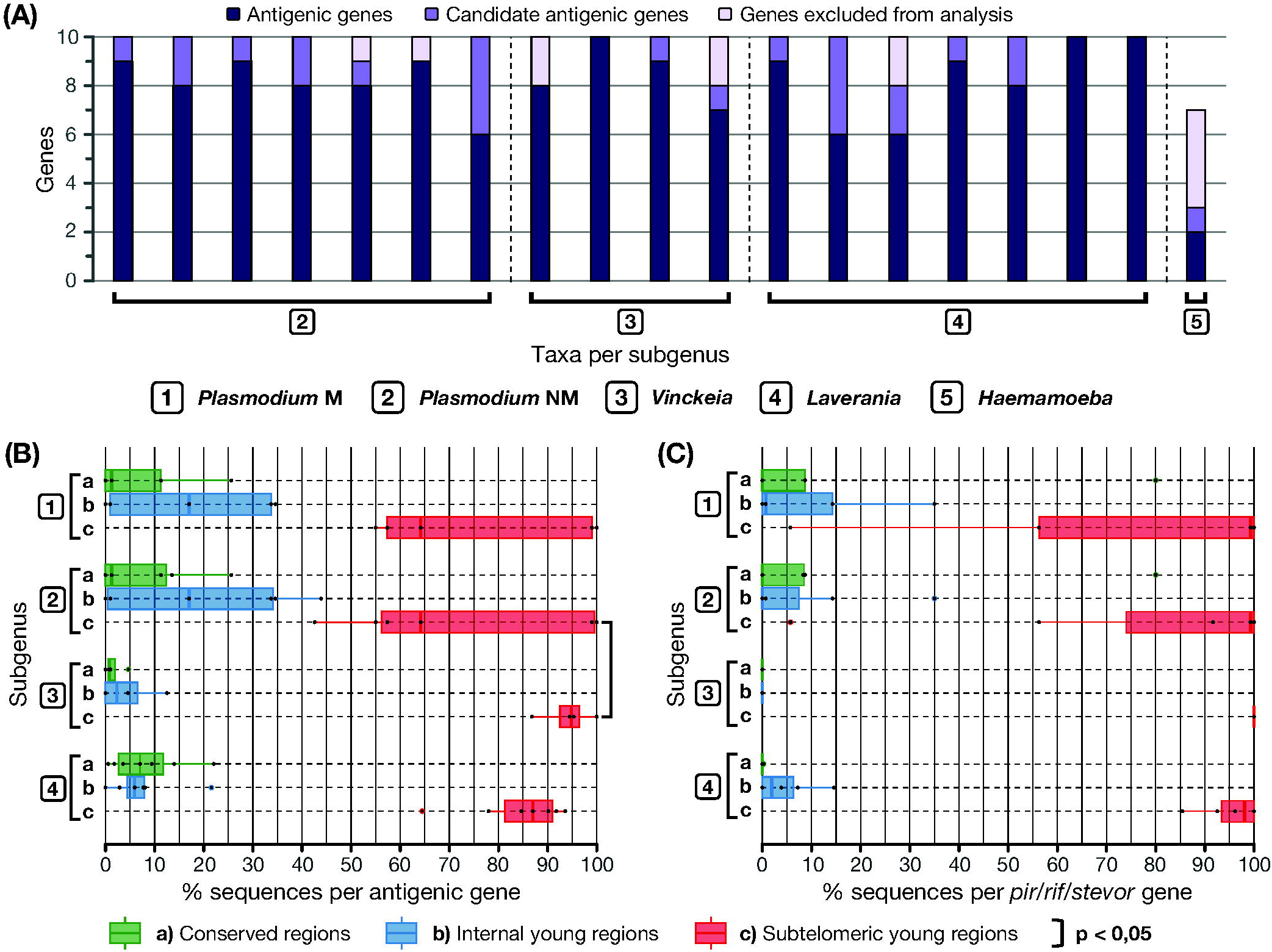
Analysis of antigenic genes distribution on chromosomes. **(A)** Classification of predominant genes in young regions according to the literature (Supplementary material). **(B)** Average percentage of sequences per antigenic gene family in each chromosome region. Genes in *Vinckeia* exhibited a significantly higher percentage of subtelomeric sequences compared to *Plasmodium* (Welch, t = 2,0594, p = 0,039). **(C)** Average percentage of sequences per *pir/rif/stevor* gene family (*pir* in *Vinckeia* and *Plasmodium, rif/stevor* in *Laverania*) in each chromosome region. A higher percentage of the sequences of these genes tend to locate preferentially in subtelomeric young regions. *Plasmodium* NM = non-monophyletic (all species), *Plasmodium* M = monophyletic (monophyletic subgroup).

The distribution of antigenic genes on chromosome maps (Fig. 3; for the full maps see Supplementary material) revealed a tendency for these genes to prefer subtelomeric regions (Fig. 4B). *Vinckeia* species exhibit the highest averages (>85%) of the number of sequences per gene family in subtelomeric young regions, and thus, a low variation is observed in the distribution of this trait. As a result, this subgenus exhibits a significantly higher average of this trait than *Plasmodium* (Welch, t = 2.0594, p = 0.039), but not significantly more than *Laverania* (T-Student, t = 1.7842, p = 0.054). In contrast, *Plasmodium* is the subgenus (even including only the monofiletic subgroup) that shows the highest variation in the averages of the number of sequences per gene family in the different chromosomal regions, while *Laverania* shows an intermediate pattern of variation (Fig. 4B). In the case of *Haemamoeba* (*P. relictum*), there is no clear location preference in the few antigenic genes detected.

All *Plasmodium* and *Vinckeia* species sampled presented *pir* genes, while 86% of *Laverania* species exhibited *rif/stevor* genes. When analyzing the distribution of sequences from these families, it was found that they tend to be located preferentially in the subtelomeres (Fig. 4C). This preference is most evident in *Vinckeia* where all the species exhibit a consistent pattern of having *pir* sequences in subtelomeric regions. In contrast, *Plasmodium* shows a high variation in the average percentage of *pir* sequences in each chromosomal region; and *Laverania* shows an intermediate variation for its *rif/stevor* sequences.

### 4.4 The intensity of ectopic recombination of subtelomeres to produce antigenic diversity

Analysis of the distribution of the CERAD regions shows a tendency to concentrate these regions in the subtelomeres, and not in internal regions, in the *Vinckeia, Laverania* and *Plasmodium* groups (Fig. 5). This tendency is less clear in *Laverania* than in *Vinckeia*, and even less clear in *Plasmodium* (the complete group and the monophyletic subgroup) where there is a high variation of this pattern among its species. Also, *Vinckeia* showed a higher distribution of CERAD regions in subtelomeres than *Laverania* (T-Student, t = 2.6121, p = 0.0141) and *Plasmodium* (Wilcoxon-Mann-Whitney, W = 25, p = 0.0224), and a significantly lower distribution of the percentage of chromosomes with internal CERAD regions than *Laverania* (T-Student, t = -1.993, p = 0.0387). On the other hand, no CERAD regions were detected in *Haemamoeba* (*P. relictum*), most likely due to the few antigenic genes and young regions found.

**Figure 5.**
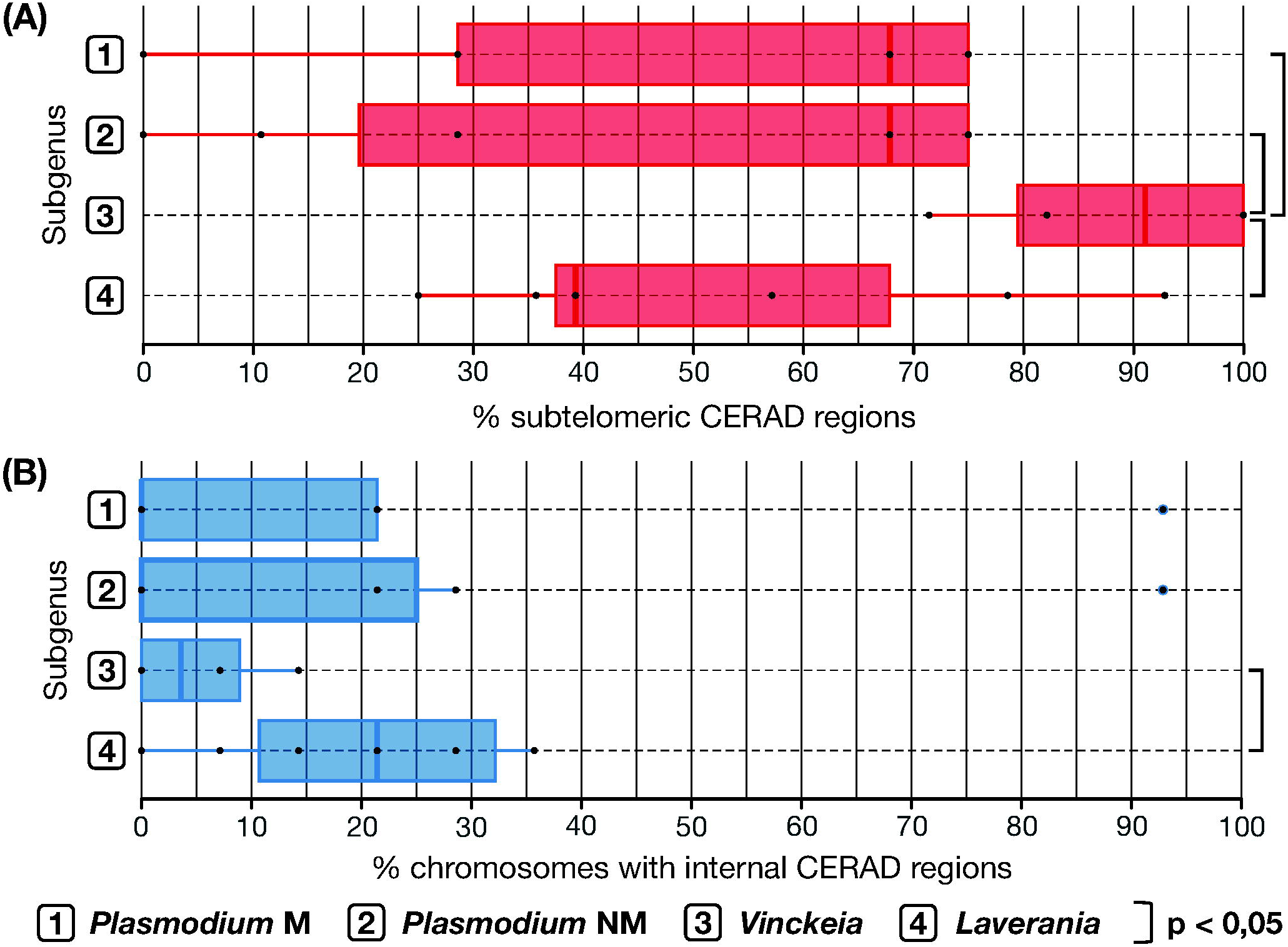
Analysis of presence of chromosomal CERAD regions. **(A)** Percentage of subtelomeric CERAD regions. *Vinckeia* has significantly higher percentages than *Laverania* (T-Student, t = 2,6121, p = 0,0141), *Plasmodium* NM (Wilcoxon-Mann-Whitney, W = 25, p = 0,0224), and *Plasmodium* M (T-Student, t = 2.1551, df = 7, p = 0.03405). **(B)** Percentage of chromosomes with internal CERAD regions. *Vinckeia* has significantly lower percentages than *Laverania* (T-Student, t = -1,993, p = 0,0387). CERAD regions = Candidate regions to undergo Ectopic Recombination to generate Antigenic Diversity, *Plasmodium* NM = non-monophyletic (all species), *Plasmodium* M = monophyletic (monophyletic subgroup).

The evaluation of the intensity of recombination to produce antigenic variation across the phylogeny shows a greater intensity of this mechanism in *Vinckeia*, an intermediate intensity in *Laverania*, high variation in *Plasmodium*, and zero intensity in *Haemamoeba* (Fig. 6A). These results are consistent with the presence of antigenic genes, particularly *pir/rif/stevor* (Fig. 6B-C), and the distribution of CERAD regions on chromosomes (Fig. 6D-E). Taken together, these features mark a distinct pattern in each subgenus. In *Vinckeia*, this pattern is characterized by a high intensity of this mechanism to produce antigenic diversity, an accumulation of CERAD regions in subtelomeres rather than in internal parts of the chromosomes, and a high number of *pir* genes. Meanwhile, in *Laverania*, an intermediate level of this mechanism is observed, which gradually increases as it approaches the *P. falciparum* and *P. praefalciparum* clade, occurring in conjunction with the increase in the percentage of CERAD regions and the number of *rif/stevor* genes. On the other hand, *Plasmodium* shows abrupt changes in the intensity levels of the mechanism and in the other evaluated characteristics, thus exhibiting the high variation that characterizes this subgenus, which is also present in its monophyletic subgroup.

**Figure 6.**
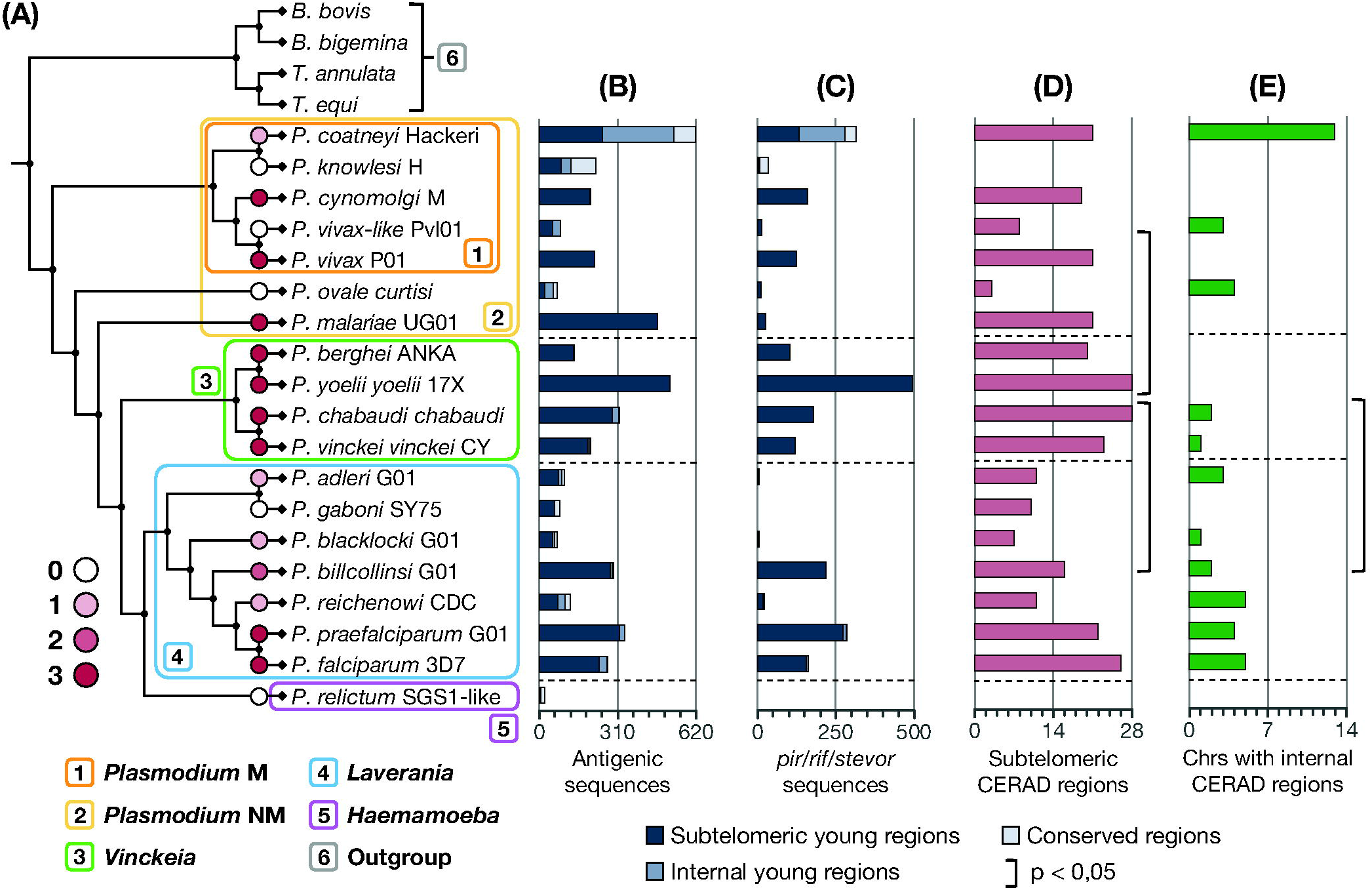
Comparison of genomic characteristics associated with ectopic recombination of subtelomeres to generate antigenic variation in *Plasmodium* species. **(A)** Intensity levels of ectopic recombination to produce antigenic diversity placed in the phylogeny of *Plasmodium*. **(B)** Number of antigenic sequences in each chromosomal region. **(C)** Number of *pir/rif/stevor* sequences (*pi*r in *Vinckeia* and Plasmodium, *rif/stevor* in *Laverania*) in each chromosomal region. **(D)** Number of CERAD subtelomeres per species. *Vinckeia* exhibits a significantly higher number of CERAD subtelomeric regions than *Laverania* (T-Student, t = 2,612, p = 0,0141) and *Plasmodium* (Wilcoxon-Mann-Whitney, W = 25, p = 0,0224). **(E)** Number of chromosomes with internal CERAD regions per species. *Vinckeia* has a significantly lower number of chromosomes with internal CERAD regions than *Laverania* (T-Student, t = -1,993, p = 0,0387). CERAD regions = Candidate regions to undergo Ectopic Recombination to generate Antigenic Diversity, *Plasmodium* NM = non-monophyletic (all species), *Plasmodium* M = monophyletic (monophyletic subgroup).

## 5 Discussion

This study shows for the first time a distribution of the phenomenon of ectopic recombination of subtelomeres for the generation of antigenic diversity in the genus *Plasmodium*, represented by 19 species of the subgenera *Plasmodium, Vinckeia, Laverania* and *Haemamoeba*. This mechanism, previously described only in *P. falciparum* with cytogenetic (Freitas-Junior, *et al*., 2000) and phylogenomic (Cerón-Romero, *et al*., 2018) data, can be inferred in the different clades by analyzing the distribution of genomic features expected from it such as a high presence of recombination and a high concentration of antigenic genes towards subtelomeres (Fig. 6). In addition, the contrast of the presence of these characteristics with the phylogeny of the group allows us to establish hypotheses about the origin and evolution of this molecular mechanism to generate antigenic diversity. Based on this, the results of this work provide three important findings: 1) The phylogeny of *Plasmodium* does not support the subgenus *Plasmodium* as monophyletic; 2) Regardless of the discordance of the phylogeny in this study and others previously published (Galen, *et al*., 2018; Pacheco, *et al*., 2018; Escalante, *et al*., 2022), *Vinckeia* shows a consistent pattern of high levels of intensity of this molecular mechanism in all its species, whereas *Laverania* exhibits a pattern of intermediate intensity and *Plasmodium* shows high variation (from zero to high intensity); 3) This molecular mechanism has been evolutionarily more associated with *pir* and *rif/stevor* genes, which fuels the debate about the homology of these gene families (Cunningham, *et al*., 2010; Harrison, *et al*., 2020).

The phylogeny reconstructed in this study (Fig. 1) contrasts with the most accepted proposal on the phylogeny of the genus *Plasmodium*, in which the subgenus *Plasmodium* is monophyletic, and bird and reptile parasites are the first lineages to diverge (i.e., closer to the root). However, it is important to keep in mind that this proposal arose from early studies based on analyses with mitochondrial DNA and/or few nuclear loci (Escalante, *et al*., 1998; Perkins & Schall, 2002; Pacheco, *et al*., 2018; Martinsen, *et al*., 2008; Krief, *et al*., 2010). More recent studies with genomic data have reported mixed results. Some studies support the monophyly of *Plasmodium* (Hayakawa, *et al*., 2008; Pacheco, *et al*., 2011; Loy, *et al*., 2017; Escalante, *et al*., 2022), while others reject it (Rutledge, *et al*., 2017; Bo□hme, *et al*., 2018). The pattern observed in this study with the subgenus *Plasmodium* at the base of the phylogeny was also obtained in a recent study, but it was interpreted as a phylogenetic artifact because of the attraction between this subgenus and the outgroup due to their similarity in GC content (Galen, *et al*., 2018). The authors tried to demonstrate this effect by generating trees after removing the “base composition bias”. However, their criterion of compositional bias is highly debatable because the GC content, which only varies between 24.5% and 43.7%, was determined from a database of only 21 genes, and because they are only considering a scenario in which the slightly higher GC content of *Plasmodium* is a derived trait.

The lack of consensus in studies with genomic data may be largely due to the difference in the number of sampled species, the lack of contrast of different phylogenetic approaches, and assuming a root for the phylogeny rather than inferring it. This has been evidenced in studies that, for example, root the *Plasmodium* tree using the clade that infects birds and/or reptiles (Hayakawa, *et al*., 2008; Pacheco, *et al*., 2011; Loy, *et al*., 2017; Escalante, *et al*., 2022)(Hayakawa, *et al*., 2008), employ three or fewer methods of phylogenetic inference (Pacheco, *et al*., 2011; Martinsen, *et al*., 2008; Krief, *et al*., 2010), or include a large number of species in the analysis, but the genetic material used is reduced to fewer than 30 nuclear and/or mitochondrial genes (Martinsen, *et al*., 2008; Pacheco, *et al*., 2018; Galen, *et al*., 2018; Krief, *et al*., 2010). Considering these three reasons, the phylogenetic analysis performed in this study is the most robust to date. However, we recognized that the results we obtained may vary significantly with the inclusion of more taxa, particularly related to *Haemamoeba, P. ovale* and *P. malariae*. Therefore, the interpretations made for the rest of the analyses were done considering different evolutionary scenarios (e.g., *Plasmodium* as a monophyletic and non-monophyletic group).

Ectopic recombination of subtelomeres to produce antigenic diversity shows different levels of significance in *Vinckeia, Laverania, Haemamoeba* and *Plasmodium*, proving to be clade-specific. Our results show that *Vinckeia* is the subgenus with the most uniform pattern among species (Figs. 2-6), characterized by the presence of ectopic recombination of subtelomeres at high levels, suggesting that this feature may have been crucial for the evolution of this group. In the case of *Laverania*, this mechanism seems to be important but no more than in *Vinckeia* (intermediate intensity) and becomes more important as one progresses towards the *P. falciparum* and *P. praefalciparum* clade (Fig. 6).

Consistent with this intermediate intensity in *Laverania* and in contrast to what was observed in *Vinckeia*, the results obtained suggest that in some cases, internal chromosomal regions of *Laverania* may ectopically recombine with the subtelomeres, as has been proposed for *var* genes in *P. falciparum* (Marty, *et al*., 2006; Claessens, *et al*., 2014). In the case of *P. relictum* (of the *Haemamoeba* clade), this mechanism does not seem to be important to promote antigenic diversity (Figs. 3 and 6). If present, this mechanism could have acquired another function, then antigenic diversity is promoted by other means (Pain, *et al*., 2008; Zhang, *et al*., 2019). On the other hand, the subgenus *Plasmodium* shows a high variation of patterns across species, suggesting that this mechanism is important for only half of them (Fig. 6). Such variation is shown even when excluding *P. ovale* and *P. malariae* and focusing only on the remaining taxa that form the monophyletic clade. This suggests that in *Plasmodium*, the importance of this mechanism was lost or acquired several times independently.

Considering the differences among the subgenera in their patterns of the intensity of ectopic recombination to generate antigenic diversity, different evolutionary scenarios can be proposed to explain the importance of this mechanism for each of them. Based on the phylogeny of this study (Figs. 1A and 6), we can infer that this mechanism appeared and became important independently on several occasions in *Plasmodium*, whereas two scenarios may have occurred in the *Vinckeia-Laverania-Haemamoeba* clade. The first scenario is that an independent acquisition occurred in the ancestors of *Vinckeia* and *Laverania*, with different levels of importance in both clades. The other scenario is that the acquisition of this mechanism occurred in the ancestor of the *Vinckeia-Laverania-Haemamoeba* clade, with independent loss in *Haemamoeba* and one *Laverania* species. The degree of probability of both scenarios depends largely on whether future studies evidence this mechanism in other *Haemamoeba* species. On the other hand, according to the phylogeny with the avian clades as the earliest divergent groups (Galen, *et al*., 2018; Escalante, *et al*., 2022), the most parsimonious scenario is that this trait appeared after the divergence of the avian groups, with different consequences for each clade: intermediate and gradual importance in *Laverania*, absolute importance in *Vinckeia*, and independent losses in *Plasmodium*. However, if other *Haemamoeba* species have this trait, it can also be an ancestral trait of the four subgenera with multiple independent losses.

The mechanism of ectopic recombination of subtelomeres is more linked to the generation of diversity of *pir* and *rif/stevor* gene families than to other gene families, reviving the debate as to whether these families are part of the same superfamily. Although studies based on the comparison of protein structures, which are more conserved and useful to detect homology than sequences, have determined significant differences between PIR and RIF/STEVOR (Harrison, *et al*., 2020), the values to establish significant differences can be arbitrary and debatable, especially when talking about proteins with a high evolutionary rate, as in this case (Claessens, *et al*., 2014; Rich & Ayala, 2000; Hernandez-Rivas, *et al*., 1996). In fact, according to our analysis, *rif* and *pir* are among the most recombinant gene families (Table S4). Therefore, the results obtained suggest one more feature in common between these proteins that may contribute to future studies aimed to establish homology among them. But at the same time, further work to clarify whether there is homology among these proteins would be useful to establish whether the association between the mechanism of ectopic recombination of subtelomeres and the diversity of these gene families is ancestral in nature.

In conclusion, we can infer from this study that ectopic recombination of subtelomeres is the main mechanism for generating diversity in the *pir* and *rif/stevor* genes, which explains the difference in the intensity of this mechanism in different *Plasmodium* clades. However, it is important to mention some of the limitations that we encountered during the execution of the analyses. For example, although this study improves several aspects of previous phylogenetic studies, we recognized that the inclusion of new taxa, especially from *Haemamoeba*, could alter the phylogenetic topology proposed here. Anticipating this limitation, the evolutionary scenarios we discussed also consider alternate phylogenetic topologies. Likewise, one could argue that the variation in the intensity of this molecular mechanism in the *Plasmodium, Laverania*, and *Vinckeia* subgenera depends on the number of sampled species. However, the high variation described for *Plasmodium* also applies to its monophyletic subgroup (Figs. 2D, 4B-C, 5 and 6), and the variation in the intensity of this mechanism in *Laverania*, can be better explained by an evolutionary pattern of a gradual increase of the importance of this mechanism towards a part of its phylogeny. Finally, it is worth noting that the inferences made here about the presence of this molecular mechanism depend on the expected consequences of this mechanism in the genome, such as the presence of subtelomeric young regions with a high density of antigenic genes. Therefore, future studies focused on performing analyses on the presence of the protein machinery, still unknown, that carries out this process at the cellular level (Figueiredo & Scherf, 2005; Hernández-Rivas, *et al*., 2013) would be key to validating the proposals presented in this study.

## 7 Author Contributions

M.A.C.R and C.M.E conceived of the study and broad approach, and designed the experiments in collaboration with H.C.H, C.M.E performed the analyses. C.M.E and M.A.C.R wrote the manuscript with input from H.C.H.

## 8 Acknowledgments

We thank the Scientific Computing Laboratory (LACCo) of CIBioFi for providing computational resources and, the Department of Biology of the Universidad del Valle for their support and suggestions during the early stages of the project.

## 10 Supplementary Material

Supplementary material can be found in figshare (https://doi.org/10.6084/m9.figshare.22179071.v7)

## 11 Data Availability Statement

Raw and analyzed data are deposited in OSF (https://doi.org/10.17605/OSF.IO/NFRG4)

## Reference styles

Al-Khedery, B., Barnwell, J.W. & Galinski, M.R., 1999. Antigenic Variation in Malaria: a 3′ Genomic Alteration Associated with the Expression of a P. knowlesi Variant Antigen. Molecular Cell, 3(2), pp.131–141. https://doi.org/10.1016/S1097-2765(00)80304-4.

Arkhipova, I.R. & Morrison, H.G., 2001. Three retrotransposon families in the genome of Giardia lamblia: Two telomeric, one dead. Proceedings of the National Academy of Sciences, 98(25), pp.14497–14502. https://doi.org/10.1073/pnas.231494798.

Aunin, E., et al., 2020. Genomic and transcriptomic evidence for descent from Plasmodium and loss of blood schizogony in Hepatocystis parasites from naturally infected red colobus monkeys. PLOS Pathogens, 16(8), p.e1008717. https://doi.org/10.1371/journal.ppat.1008717.

Barry, J.D., Ginger, M.L., Burton, P. & McCulloch, R., 2003. Why are parasite contingency genes often associated with telomeres? International Journal for Parasitology, 33(1), pp.29–45. https://doi.org/10.1016/S0020-7519(02)00247-3.

Böhme, U., et al., 2018. Complete avian malaria parasite genomes reveal features associated with lineage-specific evolution in birds and mammals. Genome Research, 28(4), pp.547–560. https://doi.org/10.1101/gr.218123.116.

Borner, J., et al., 2016. Phylogeny of haemosporidian blood parasites revealed by a multi-gene approach. Molecular Phylogenetics and Evolution, 94, pp.221–231. https://doi.org/10.1016/j.ympev.2015.09.003.

Bowman, S., et al., 1999. The complete nucleotide sequence of chromosome 3 of Plasmodium falciparum. Nature, 400(6744), pp.532–538. https://doi.org/10.1038/22964.

Carlton, J.M., et al., 2008. Comparative genomics of the neglected human malaria parasite Plasmodium vivax. Nature, 455(7214), pp.757–763. https://doi.org/10.1038/nature07327.

Carlton, J.M.-R., Galinski, M.R., Barnwell, J.W. & Dame, J.B., 1999. Karyotype and synteny among the chromosomes of all four species of human malaria parasite. Molecular and Biochemical Parasitology, 101(1), pp.23–32. https://doi.org/10.1016/S0166-6851(99)00045-6.

Cerón-Romero, M.A., Nwaka, E., Owoade, Z. & Katz, L.A., 2018. PhyloChromoMap, a Tool for Mapping Phylogenomic History along Chromosomes, Reveals the Dynamic Nature of Karyotype Evolution in Plasmodium falciparum. Genome Biology and Evolution, 10(2), pp.553–561. https://doi.org/10.1093/gbe/evy017.

Cheng, Q., et al., 1998. stevor and rif are Plasmodium falciparum multicopy gene families which potentially encode variant antigens. Molecular and Biochemical Parasitology, 97(1), pp.161–176. https://doi.org/10.1016/S0166-6851(98)00144-3.

Claessens, A., et al., 2014. Generation of Antigenic Diversity in Plasmodium falciparum by Structured Rearrangement of Var Genes During Mitosis. PLOS Genetics, 10(12), p.e1004812. https://doi.org/10.1371/journal.pgen.1004812.

Cunningham, D., et al., 2010. The pir multigene family of Plasmodium: Antigenic variation and beyond. Molecular and Biochemical Parasitology, 170(2), pp.65–73. https://doi.org/10.1016/j.molbiopara.2009.12.010.

Emms, D.M. & Kelly, S., 2015. OrthoFinder: solving fundamental biases in whole genome comparisons dramatically improves orthogroup inference accuracy. Genome Biology, 16(1), p.157. https://doi.org/10.1186/s13059-015-0721-2.

Escalante, A.A., Cepeda, A.S. & Pacheco, M.A., 2022. Why Plasmodium vivax and Plasmodium falciparum are so different? A tale of two clades and their species diversities. Malaria Journal, 21(1), p.139. https://doi.org/10.1186/s12936-022-04130-9.

Escalante, A.A., Freeland, D.E., Collins, W.E. & Lal, A.A., 1998. The evolution of primate malaria parasites based on the gene encoding cytochrome b from the linear mitochondrial genome. Proceedings of the National Academy of Sciences, 95(14), pp.8124–8129. https://doi.org/10.1073/pnas.95.14.8124.

Figueiredo, L. & Scherf, A., 2005. Plasmodium telomeres and telomerase: the usual actors in an unusual scenario. Chromosome Research, 13(5), pp.517–524. https://doi.org/10.1007/s10577-005-0996-3.

Figueiredo, L.M., et al., 2002. A central role for Plasmodium falciparum subtelomeric regions in spatial positioning and telomere length regulation. The EMBO Journal, 21(4), pp.815–824. https://doi.org/10.1093/emboj/21.4.815.

Frank, M., et al., 2008. Frequent recombination events generate diversity within the multi-copy variant antigen gene families of Plasmodium falciparum. International journal for parasitology, 38(10), pp.1099–1109. https://doi.org/10.1016/j.ijpara.2008.01.010.

Freitas-Junior, L.H., et al., 2000. Frequent ectopic recombination of virulence factor genes in telomeric chromosome clusters of P. falciparum. Nature, 407(6807), pp.1018–1022. https://doi.org/10.1038/35039531.

Galen, S.C., et al., 2018. The polyphyly of Plasmodium: comprehensive phylogenetic analyses of the malaria parasites (order Haemosporida) reveal widespread taxonomic conflict. Royal Society Open Science, 5(5), p.171780. https://doi.org/10.1098/rsos.171780.

Gardner, M.J., et al., 2002. Genome sequence of the human malaria parasite Plasmodium falciparum. Nature, 419(6906), pp.498–511. https://doi.org/10.1038/nature01097.

Guindon, S., et al., 2010. New Algorithms and Methods to Estimate Maximum-Likelihood Phylogenies: Assessing the Performance of PhyML 3.0. Systematic Biology, 59(3), pp.307–321. https://doi.org/10.1093/sysbio/syq010.

Harrison, T.E., et al., 2020. Structure of the Plasmodium-interspersed repeat proteins of the malaria parasite. Proceedings of the National Academy of Sciences, 117(50), pp.32098–32104. https://doi.org/10.1073/pnas.2016775117.

Hayakawa, T., et al., 2008. Big Bang in the Evolution of Extant Malaria Parasites. Molecular Biology and Evolution, 25(10), pp.2233–2239. https://doi.org/10.1093/molbev/msn171.

Hernández-Rivas, R., et al., 2013. Impact of chromosome ends on the biology and virulence of Plasmodium falciparum. Molecular and Biochemical Parasitology, 187(2), pp.121–128. https://doi.org/10.1016/j.molbiopara.2013.01.003.

Hernandez-Rivas, R., Hinterberg, K. & Scherf, A., 1996. Compartmentalization of genes coding for immunodominant antigens to fragile chromosome ends leads to dispersed subtelomeric gene families and rapid gene evolution in Plasmodium falciparum. Molecular and Biochemical Parasitology, 78(1), pp.137–148. https://doi.org/10.1016/S0166-6851(96)02618-7.

Hoeijmakers, W.A.M., et al., 2012. Plasmodium falciparum centromeres display a unique epigenetic makeup and cluster prior to and during schizogony. Cellular Microbiology, 14(9), pp.1391–1401. https://doi.org/10.1111/j.1462-5822.2012.01803.x.

Janssen, C.S., Barrett, M.P., Turner, C.M.R. & Phillips, R.S., 2002. A large gene family for putative variant antigens shared by human and rodent malaria parasites. Proceedings of the Royal Society of London. Series B: Biological Sciences, 269(1489), pp.431–436. https://doi.org/10.1098/rspb.2001.1903.

Janssen, C.S., Phillips, R.S., Turner, C.M.R. & Barrett, M.P., 2004. Plasmodium interspersed repeats: the major multigene superfamily of malaria parasites. Nucleic Acids Research, 32(19), pp.5712–5720. https://doi.org/10.1093/nar/gkh907.

Katoh, K., Kuma, K., Toh, H. & Miyata, T., 2005. MAFFT version 5: improvement in accuracy of multiple sequence alignment. Nucleic Acids Research, 33(2), pp.511–518. https://doi.org/10.1093/nar/gki198.

Kemp, D.J., Thompson, J.K., Walliker, D. & Corcoran, L.M., 1987. Molecular karyotype of Plasmodium falciparum: conserved linkage groups and expendable histidine-rich protein genes. Proceedings of the National Academy of Sciences, 84(21), pp.7672–7676. https://doi.org/10.1073/pnas.84.21.7672.

Kooij, T.W.A., et al., 2005. A Plasmodium Whole-Genome Synteny Map: Indels and Synteny Breakpoints as Foci for Species-Specific Genes. PLOS Pathogens, 1(4), p.e44. https://doi.org/10.1371/journal.ppat.0010044.

Krief, S., et al., 2010. On the Diversity of Malaria Parasites in African Apes and the Origin of Plasmodium falciparum from Bonobos. PLOS Pathogens, 6(2), p.e1000765. https://doi.org/10.1371/journal.ppat.1000765.

Little, T.S., et al., 2021. Analysis of pir gene expression across the Plasmodium life cycle. Malaria Journal, 20(1), p.445. https://doi.org/10.1186/s12936-021-03979-6.

Loy, D.E., et al., 2017. Out of Africa: origins and evolution of the human malaria parasites Plasmodium falciparum and Plasmodium vivax. International journal for parasitology, 47(2–3), pp.87–97. https://doi.org/10.1016/j.ijpara.2016.05.008.

Martinsen, E.S., Perkins, S.L. & Schall, J.J., 2008. A three-genome phylogeny of malaria parasites (Plasmodium and closely related genera): Evolution of life-history traits and host switches. Molecular Phylogenetics and Evolution, 47(1), pp.261–273. https://doi.org/10.1016/j.ympev.2007.11.012.

Marty, A.J., et al., 2006. Evidence that Plasmodium falciparum chromosome end clusters are cross-linked by protein and are the sites of both virulence gene silencing and activation. Molecular Microbiology, 62(1), pp.72–83. https://doi.org/10.1111/j.1365-2958.2006.05364.x.

Minh, B.Q., Nguyen, M.A.T. & von Haeseler, A., 2013. Ultrafast Approximation for Phylogenetic Bootstrap. Molecular Biology and Evolution, 30(5), pp.1188–1195. https://doi.org/10.1093/molbev/mst024.

Morel, B., et al., 2022. SpeciesRax: A Tool for Maximum Likelihood Species Tree Inference from Gene Family Trees under Duplication, Transfer, and Loss. Molecular Biology and Evolution, 39(2), p.msab365. https://doi.org/10.1093/molbev/msab365.

Nguyen, L.-T., Schmidt, H.A., von Haeseler, A. & Minh, B.Q., 2015. IQ-TREE: A Fast and Effective Stochastic Algorithm for Estimating Maximum-Likelihood Phylogenies. Molecular Biology and Evolution, 32(1), pp.268–274. https://doi.org/10.1093/molbev/msu300.

Otto, T.D., et al., 2018. Long read assemblies of geographically dispersed Plasmodium falciparum isolates reveal highly structured subtelomeres. Wellcome Open Research, 3, p.52. https://doi.org/10.12688/wellcomeopenres.14571.1.

Pacheco, M.A., et al., 2011. Timing the origin of human malarias: the lemur puzzle. BMC Evolutionary Biology, 11(1), p.299. https://doi.org/10.1186/1471-2148-11-299.

Pacheco, M.A., et al., 2018. Mode and Rate of Evolution of Haemosporidian Mitochondrial Genomes: Timing the Radiation of Avian Parasites. Molecular Biology and Evolution, 35(2), pp.383–403. https://doi.org/10.1093/molbev/msx285.

Pain, A., et al., 2008. The genome of the simian and human malaria parasite Plasmodium knowlesi. Nature, 455(7214), pp.799–803. https://doi.org/10.1038/nature07306.

Perkins, S.L., 2014. Malaria’s Many Mates: Past, Present, and Future of the Systematics of the Order Haemosporida. Journal of Parasitology, 100(1), pp.11–25. https://doi.org/10.1645/13-362.1.

Perkins, S.L. & Schall, J., 2002. A molecular phylogeny of malarial parasites recovered from cytochrome b gene sequences. Journal of Parasitology, 88(5), pp.972–978. https://doi.org/10.1645/0022-3395(2002)088[0972:AMPOMP]2.0.CO;2.

del Portillo, H.A., et al., 2001. A superfamily of variant genes encoded in the subtelomeric region of Plasmodium vivax. Nature, 410(6830), pp.839–842. https://doi.org/10.1038/35071118.

de Queiroz, A. & Gatesy, J., 2007. The supermatrix approach to systematics. Trends in Ecology & Evolution, 22(1), pp.34–41. https://doi.org/10.1016/j.tree.2006.10.002.

Rabiee, M., Sayyari, E. & Mirarab, S., 2019. Multi-allele species reconstruction using ASTRAL. Molecular Phylogenetics and Evolution, 130, pp.286–296. https://doi.org/10.1016/j.ympev.2018.10.033.

Reed, J., et al., 2021. Telomere length dynamics in response to DNA damage in malaria parasites. iScience, [online] 24(2). https://doi.org/10.1016/j.isci.2021.102082.

Rich, S.M. & Ayala, F.J., 2000. Population structure and recent evolution of Plasmodium falciparum. Proceedings of the National Academy of Sciences, 97(13), pp.6994–7001. https://doi.org/10.1073/pnas.97.13.6994.

Rich, S.M. & Xu, G., 2011. Resolving the phylogeny of malaria parasites. Proceedings of the National Academy of Sciences, 108(32), pp.12973–12974. https://doi.org/10.1073/pnas.1110141108.

Rutledge, G.G., et al., 2017. Plasmodium malariae and P. ovale genomes provide insights into malaria parasite evolution. Nature, 542(7639), pp.101–104. https://doi.org/10.1038/nature21038.

Salomaki, E.D., et al., 2021. Gregarine single-cell transcriptomics reveals differential mitochondrial remodeling and adaptation in apicomplexans. BMC Biology, 19(1), p.77. https://doi.org/10.1186/s12915-021-01007-2.

Scherf, A., Figueiredo, L.M. & Freitas-Junior, L.H., 2001. Plasmodium telomeres: a pathogen’s perspective. Current Opinion in Microbiology, 4(4), pp.409–414. https://doi.org/10.1016/S1369-5274(00)00227-7.

Sharp, P.M., Plenderleith, L.J. & Hahn, B.H., 2020. Ape Origins of Human Malaria. Annual Review of Microbiology, 74(1), pp.39–63. https://doi.org/10.1146/annurev-micro-020518-115628.

Silva Pereira, S., et al., 2020. Variant antigen diversity in Trypanosoma vivax is not driven by recombination. Nature Communications, 11(1), p.844. https://doi.org/10.1038/s41467-020-14575-8.

Stecher, G., Tamura, K. & Kumar, S., 2020. Molecular Evolutionary Genetics Analysis (MEGA) for macOS. Molecular Biology and Evolution, 37(4), pp.1237–1239. https://doi.org/10.1093/molbev/msz312.

Su, X., et al., 1995. The large diverse gene family var encodes proteins involved in cytoadherence and antigenic variation of Plasmodium falciparum-infected erythrocytes. Cell, 82(1), pp.89–100. https://doi.org/10.1016/0092-8674(95)90055-1.

Suyama, M., Torrents, D. & Bork, P., 2006. PAL2NAL: robust conversion of protein sequence alignments into the corresponding codon alignments. Nucleic Acids Research, 34(Suppl_2), pp.W609–W612. https://doi.org/10.1093/nar/gkl315.

Swofford, D., 2002. PAUP*: Phylogenetic Analysis Using Parsimony (*and Other Methods). Available at: <https://paup.phylosolutions.com/>.

WHO, 2014. World malaria report 2014. [online] World Health Organization. Available at: <https://www.who.int/publications/i/item/9789241564830> [Accessed 10 February 2023].

Willson, J., et al., 2022. DISCO: Species Tree Inference using Multicopy Gene Family Tree Decomposition. Systematic Biology, 71(3), pp.610–629. https://doi.org/10.1093/sysbio/syab070.

Zhang, C., Scornavacca, C., Molloy, E.K. & Mirarab, S., 2020. ASTRAL-Pro: Quartet-Based Species-Tree Inference despite Paralogy. Molecular Biology and Evolution, 37(11), pp.3292–3307. https://doi.org/10.1093/molbev/msaa139.

Zhang, X., et al., 2019. Rapid antigen diversification through mitotic recombination in the human malaria parasite Plasmodium falciparum. PLOS Biology, 17(5), p.e3000271. https://doi.org/10.1371/journal.pbio.3000271

