## Supplementary material for "Tracking the intensity of the mechanism to produce antigenic diversity by subtelomeric ectopic recombination across the phylogeny of *Plasmodium* parasites": Supplementary_material.docx

**Supplementary material datasets:**

1. **Dataset 1:** Database for phylogenetic reconstruction of the genus Plasmodium.
2. **Dataset 2:** Trees of the genus Plasmodium generated with six phylogenetic tools (Majority-rule consensus tree of the 6 trees obtained).
3. **Dataset 3:** Database for chromosome mapping of gene conservation.
4. **Dataset 4:** List of predominant gene families in young regions per species.
5. **Dataset 5:** Classification of gene families (with the highest number of sequences in young regions) based on the nature of its products.
6. **Dataset 6:** Chromosome maps of 19 *Plasmodium* species showing the gene conservation profile, presence of young regions, and distribution of antigenic genes.
1. *P. adleri* (Page 1)
2. *P. berghei* (Page 2)
3. *P. billcollinsi* (Page 3)
4. *P. blacklocki* (Page 4)
5. *P. chabaudi* chabaudi (Page 5)
6. *P. coatneyi* (Page 6)
7. *P. cynomolgi* (Page 7)
8. *P. falciparum* (Page 8)
9. *P. gaboni* (Page 9)
10. *P. knowlesi* (Page 10)
11. *P. malariae* (Page 11)
12. *P. ovale curtisi* (Page 12)
13. *P. praefalciparum* (Page 13)
14. *P. reichenowi* (Page 14)
15. *P. relictum* (Page 15)
16. *P. vivax-like* (Page 16)
17. *P. vivax* (Page 17)
18. *P. vinckei vinckei* (Page 18)
19. *P. yoelii yoelii* (Page 19)

**Supplementary table S1.** Criteria to define the four levels in which *Plasmodium* species are classified based on evidence of the intensity of production of antigenic diversity through ectopic recombination.

| **Criteria** | | **Level** | | | |
| --- | --- | --- | --- | --- | --- |
|  |  | 0 | 1 | 2 | 3 |
| Presence of recombinant subtelomeric regions | a. Less than 25% of chromosomal ends appear to be candidate recombinant subtelomeric young regions* b. Between 25-39% of chromosomal ends appear to be candidate recombinant subtelomeric young regions * c. Between 40-65% of chromosomal ends appear to be candidate recombinant subtelomeric young regions * d. More than 65% of chromosomal ends appear to be candidate recombinant subtelomeric young regions * | X |  |  |  |
|  |  |  | X |  |  |
|  |  |  |  | X |  |
|  |  |  |  |  | X |
| Location of gene families | a. Some gene families are located in subtelomeric and internal young regions b. A negligible number of gene families are located in both subtelomeric and internal young regions | X | X |  |  |
|  |  |  |  | X | X |
| Presence of mechanisms other than recombination | a. There are gene families that do not appear to be undergoing recombination in young subtelomeres (gene cassettes) b. There may be gene families that are not undergoing recombination in young subtelomeres, but it is almost imperceptible | X | X |  |  |
|  |  |  |  | X | X |

(*) distinctive young regions in the subtelomeres with higher density (>3) of antigenic genes

**Supplementary table S2.** Number of species present in each major clade and in their respective minor clades. Sequence nomenclature employs the two-letter abbreviation defined for each clade. The full name of each clade is indicated in parenthesis next to the abbreviation.

| **Major clade** | **Minor clade** |
| --- | --- |
| Ap (Apicomplexa) = 57 | 1. ec (Eucoccidiorida) = 12 2. eu (Eugregarines) = 3 3. ha (Haemosporida) = 34 4. pi (Piroplasmida) = 8 |
| Oa (Other Alveolates) = 14 | 1. ch (Chromerida) = 2 2. ci (Ciliophora) = 6 3. df (Dinoflagellata) = 3 4. pe (Perkinsozoa) = 3 |
| Rh (Rhizaria) = 4 | 1. ce (Cercozoa) = 1 2. im (Imbricatea) = 1 3. ex (Endomyxa) = 1 4. fo (Foraminifera) = 1 |
| St (Stramenopila) = 33 | 1. bi (Bigyra) = 6 2. pn (Perenosporales) = 9 3. sa (Saprolegniales) = 8 4. oc (Ochrophyta) = 10 |
| Ds (Discoba) = 15 | 1. eb (Eubodonida) = 1 2. ty (Trypanosomatida) = 9 3. ih (Ichthyobodonidae) = 1 4. va (Vahlkampfiidae) = 3 5. jk (Jakobida) = 1 |
| Me (Metamonada) = 11 | 1. fr (Fornicata) = 6 2. pb (Parabasalia) = 3 3. px (Preaxostyla) = 2 |

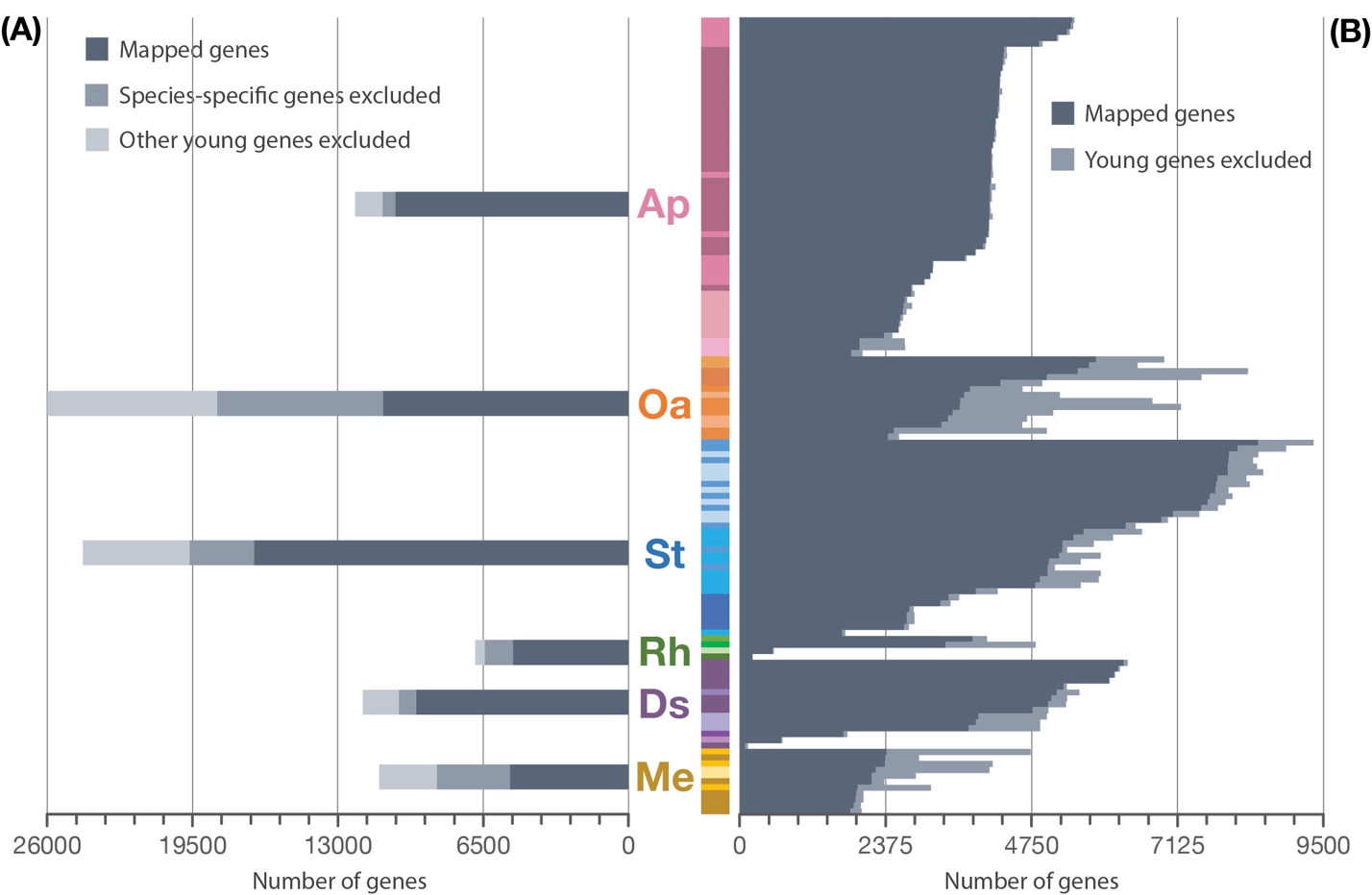

**Supplementary figure S1.** Proportion of species-specific or young genes excluded from chromosome mapping analysis. The major clades are: Apicomplexa (Ap), Other Alveolates (Oa), Stramenopila (St), Rhizaria (Rh), Discoba (Ds), Metamonada (Me) **(A)** Proportion given by major clade **(B)** Proportion given by species. The color bar indicates the diversity of minor clades present in each major clade.
