## Supplementary material for "Tracking the intensity of the mechanism to produce antigenic diversity by subtelomeric ectopic recombination across the phylogeny of *Plasmodium* parasites": Supplementary_material_dataset5.docx

**Supplementary dataset 5.** Classification of gene families (with the highest number of sequences in young regions) based on the nature of its products. To classify the gene families as antigenic, candidate or discarded, we reviewed the available literature on the product of each of them.

| **Gene Family** | **Product** | **Function** | **References** | **Classification** |
| --- | --- | --- | --- | --- |
| OG0000000 | CDPK, CaMKs | Development of sporozoites and subsequent invasion of hepatocytes; merozoite adhesion and subsequent red blood cell invasion; development of ookinetes and subsequent invasion of the midgut of the mosquito. | [Ghartey-Kwansah et al. (2020); Kato et al. (2008)](https://doi.org/10.1177/0963689719884888) | Antigenic |
| OG0000004 | Heat shock proteins (eg DnaJ) | They play an important role in the circulation and folding of proteins exported to erythrocytes, thus contributing to the pathology of malaria. | [Daniyan et al. (2016](https://doi.org/10.1371/journal.pone.0148517));  [Pesce et al. (2008)](https://doi.org/10.1016/j.biocel.2008.06.011) | Candidate |
| OG0000023 | DNA/RNA helicases (ATPases), helicases | Important role in replication and transmission of the genetic information of the parasite. | [Tuteja & Pradhan (2006)](https://doi.org/10.1016/j.gene.2006.03.007) | Discarded |
| OG0000043 | Several products: antigen (1), helicases (3), among others (3) | Role of helicases in DNA replication and transmission; Antigens promote the invasion process of parasites. | [Tuteja & Pradhan (2006)](https://doi.org/10.1016/j.gene.2006.03.007);  [Fan et al. (2020)](https://doi.org/10.3389/fpubh.2020.00148) | Discarded |
| OG0000065 | Acyl-CoA synthetases | Elevated levels in infected erythrocytes suggests its importance in the early development of the parasite within the erythrocyte and in the schizogony or invasion process. | [Matesanz et al. (2003](https://doi.org/10.1016/S0166-6851(02)00242-6));  [Prata](https://doi.org/10.1021/acsinfecdis.1c00414) et al. (2021) | Antigenic |
| OG0000203 | Lysophospholipase | Important for parasite development within erythrocytes and crucial for normal schizogonic division. | [Sheokand et al. 2021](https://www.biorxiv.org/content/10.1101/2021.07.01.450682v1);  [Asad et al. (2021)](https://doi.org/10.1186/s12915-021-01042-z) | Candidate |
| OG0000468 | Tryptophan-rich antigen (TRAgs) | Associated with the membrane of infected erythrocytes, they can bind to the surface of uninfected erythrocytes, triggering an immune response. | [Wang et al. (2015](https://doi.org/10.1128/IAI.03067-14));  [Burns et al. (2000)](https://doi.org/10.1016/S0166-6851(00)00252-8) | Antigenic |
| OG0000547 | Serine/threonine protein kinases (FIKK gene family) | They regulate the rigidity and cytoadhesion of infected erythrocytes, thus contributing to the survival of the parasite. | [Davies et al. (2020](https://doi.org/10.1038/s41564-020-0702-4));  [Nunes et al. (2010)](https://doi.org/10.1371/journal.pone.0011747) | Antigenic |
| OG0000722 | PIR protein | Increased expression in stages that invade and reside in erythrocytes suggests an important role in parasite biology. | [Little et al. (2021](https://doi.org/10.1186/s12936-021-03979-6));  [Janssen et al. (2004);](https://doi.org/10.1093/nar/gkh907) [Otto et al. (2014)](https://doi.org/10.1186/s12915-014-0086-0) | Antigenic |
| OG0000730 | BE | Antigen present in infected erythrocytes, during the trophozoite and schizont stage. Important for survival and successful infection by the parasite. | [Aoki et al. (2002](https://doi.org/10.1074/jbc.M207145200));  [Arisue et al. (2020)](https://doi.org/10.1186/s13071-020-04044-y) | Antigenic |
| OG0000751 | DnaJ protein, protein with PHIST domain or RESA type | Important for the survival of the parasite and the infection process. | [Pei et al. (2007](https://doi.org/10.1182/blood-2007-02-076919)); [Daniyan et al. (2016);](https://doi.org/10.1371/journal.pone.0148517)  [Yang et al. (2020)](https://doi.org/10.3389/fmicb.2020.611190) | Candidate |
| OG0000871 | Fam-a protein | Important for the invasion of erythrocytes, expressed in merozoites, keys for the development of the parasite. | [Alam et al. (2016](https://doi.org/10.1016/j.bbrc.2016.08.096));  [Zeeshan et al. (2015)](https://doi.org/10.1093/infdis/jiu558) | Antigenic |
| OG0000885 | Fam-a protein | Important for the invasion of erythrocytes, expressed in merozoites, keys for the development of the parasite. | [Alam et al. (2016](https://doi.org/10.1016/j.bbrc.2016.08.096));  [Zeeshan et al. (2015)](https://doi.org/10.1093/infdis/jiu558) | Antigenic |
| OG0001027 | RIFIN | Antigen expressed on the surface of infected erythrocytes. Evidence suggests that it is important for evading the immune system. | [Bachmann et al. (2015);](https://doi.org/10.1186/s12936-015-0784-2)  [Saito et al. (2017);](https://doi.org/10.1038/nature24994)  [Goel et al. (2015)](https://doi.org/10.1038/nm.3812) | Antigenic |
| **Gene Family** | **Product** | **Function** | **References** | **Classification** |
| OG0001028 | PIR protein | Increased expression in stages that invade and reside in erythrocytes suggests an important role in parasite biology. | [Little et al. (2021](https://doi.org/10.1186/s12936-021-03979-6));  [Janssen et al. (2004);](https://doi.org/10.1093/nar/gkh907) [Otto et al. (2014)](https://doi.org/10.1186/s12915-014-0086-0) | Antigenic |
| OG0001209 | MSP7 type protein, MSP7 | Key to host invasion by participating in initial binding to the red blood cell membrane. | [Castillo et al. (2017](https://doi.org/10.1016/j.meegid.2017.01.024));  [Kadekoppala et al. (2008)](https://doi.org/10.1128/EC.00274-08) | Antigenic |
| OG0001220 | SICA antigen, SICA-type | Produced during the trophozoite stage and then inserted into the erythrocyte membrane and exposed to the outside of the host cell. | [Korir & Galinski (2006](https://doi.org/10.1016/j.meegid.2005.01.003));  [Howard & Barnwell (1984)](https://doi.org/10.1007/978-1-4684-4571-8_5) | Antigenic |
| OG0001264 | Early transcription membrane protein | They have a role in host-parasite interactions, performing vital functions *in vivo* before infection of blood cells. | [Spielman et al. (2003](https://www.molbiolcell.org/doi/10.1091/mbc.e02-04-0240?url_ver=Z39.88-2003&rfr_id=ori:rid:crossref.org&rfr_dat=cr_pub%20%200pubmed));  [MacKellas et al. (2011);](https://doi.org/10.1111/j.1462-5822.2011.01656.x) [Garcia et al. (2009)](https://doi.org/10.1016/j.vaccine.2009.09.009) | Antigenic |
| OG0001282 | PIR protein (KIR, CYIR, KIR type, VIR7 type) | Increased expression in stages that invade and reside in erythrocytes suggests an important role in parasite biology. | [Little et al. (2021](https://doi.org/10.1186/s12936-021-03979-6));  [Janssen et al. (2004);](https://doi.org/10.1093/nar/gkh907) [Otto et al. (2014)](https://doi.org/10.1186/s12915-014-0086-0) | Antigenic |
| OG0001526 | SICAvar (type I) | Produced during the trophozoite stage and then inserted into the erythrocyte membrane and exposed to the outside of the host cell. | [Korir & Galinski (2006](https://doi.org/10.1016/j.meegid.2005.01.003));  [Howard & Barnwell (1984)](https://doi.org/10.1007/978-1-4684-4571-8_5) | Antigenic |
| OG0001643 | RIFIN | Antigen expressed on the surface of infected erythrocytes. Evidence suggests that it is important for evading the immune system. | [Bachmann et al. (2015);](https://doi.org/10.1186/s12936-015-0784-2)  [Saito et al. (2017);](https://doi.org/10.1038/nature24994)  [Goel et al. (2015)](https://doi.org/10.1038/nm.3812) | Antigenic |
| OG0001868 | *Exported Plasmodium* protein | They can induce changes in the physiology of erythrocytes important for parasite growth and survival. | [Marti & Spielmann (2013)](https://doi.org/10.1016/j.mib.2013.04.010);  [Matthews et al. (2019)](https://doi.org/10.1111/cmi.13009) | Candidate |
| OG0001869 | Fam-l protein | They appear to be exported from the parasite to blood cells. | [Rutledge et al. (2017)](https://www.nature.com/articles/nature21038) | Antigenic |
| OG0001919 | PfEMP1 | It undergoes antigenic variation to evade the immune system and changes the cytoadhesive properties of infected red blood cells so that they bind to uninfected ones. | [Pasternak & Dzikowski (2009](https://doi.org/10.1016/j.biocel.2008.12.012));  [Jensen et al. (2019)](https://doi.org/10.1111/imr.12807) | Antigenic |
| OG0001956 | *Exported Plasmodium* protein (PHIST) | PHIST family important for parasite-host interactions | [Yang et al. (2020](https://doi.org/10.3389/fmicb.2020.611190));  [Oberli et al. (2014)](https://doi.org/10.1096/fj.14-256057) | Antigenic |
| OG0002065 | Asexual protein linked to cytoadhesion | It binds to blood cell receptors. It is suggested that they are key in the adhesion to vascular endothelium receptors, a key process for the virulence of the parasite. | [Ocampo et al. (2005)](https://doi.org/10.1110/ps.04883905) | Antigenic |
| OG0002087 | Reticulocyte-binding protein (RBPs) | Invade reticulocytes (young blood cells) | [Chan et al. (2020)](https://doi.org/10.1111/cmi.13110) | Antigenic |
| OG0002136 | SICA antigen | Produced during the trophozoite stage and then inserted into the erythrocyte membrane and exposed to the outside of the host cell. | [Korir & Galinski (2006](https://doi.org/10.1016/j.meegid.2005.01.003));  [Howard & Barnwell (1984)](https://doi.org/10.1007/978-1-4684-4571-8_5) | Antigenic |
| OG0002408 | RIFIN | Antigen expressed on the surface of infected erythrocytes. Evidence suggests that it is important for evading the immune system. | [Bachmann et al. (2015);](https://doi.org/10.1186/s12936-015-0784-2)  [Saito et al. (2017);](https://doi.org/10.1038/nature24994)  [Goel et al. (2015)](https://doi.org/10.1038/nm.3812) | Antigenic |
| **Gene Family** | **Product** | **Function** | **References** | **Classification** |
| OG0002560 | PIR Protein (VIR) | Increased expression in stages that invade and reside in erythrocytes suggests an important role in parasite biology. | [Little et al. (2021](https://doi.org/10.1186/s12936-021-03979-6));  [Janssen et al. (2004);](https://doi.org/10.1093/nar/gkh907) [Otto et al. (2014)](https://doi.org/10.1186/s12915-014-0086-0) | Antigenic |
| OG0002561 | Fam-m protein | They appear to be exported from the parasite to blood cells. | [Rutledge et al. (2017)](https://www.nature.com/articles/nature21038) | Antigenic |
| OG0002744 | SURFIN | Expressed in merozoites and infected erythrocytes suggesting a role in the invasion process. | [Winter et al. (2005)](https://doi.org/10.1084/jem.20041392) | Antigenic |
| OG0002915 | RIFIN | Antigen expressed on the surface of infected erythrocytes. Evidence suggests that it is important for evading the immune system. | [Bachmann et al. (2015);](https://doi.org/10.1186/s12936-015-0784-2)  [Saito et al. (2017);](https://doi.org/10.1038/nature24994)  [Goel et al. (2015)](https://doi.org/10.1038/nm.3812) | Antigenic |
| OG0002944 | *Exported Plasmodium* proteins (PHISTa, PHISTa type, ACBP) | Important for parasite-host interactions or for parasite growth and survival. | [Yang et al. (2020](https://doi.org/10.3389/fmicb.2020.611190));  [Kumar et al. (2019);](https://doi.org/10.1021/acschembio.9b00003) [Oberli et al. (2014)](https://doi.org/10.1096/fj.14-256057) | Antigenic |
| OG0003008 | PfEMP1 | It undergoes antigenic variation to evade the immune system and changes the cytoadhesive properties of infected red blood cells so that they bind to uninfected ones. | [Pasternak & Dzikowski (2009](https://doi.org/10.1016/j.biocel.2008.12.012));  [Jensen et al. (2019)](https://doi.org/10.1111/imr.12807) | Antigenic |
| OG0003178 | PIR protein (KIR, KIR type, VIR7 type) | Increased expression in stages that invade and reside in erythrocytes suggests an important role in parasite biology. | [Little et al. (2021](https://doi.org/10.1186/s12936-021-03979-6));  [Janssen et al. (2004);](https://doi.org/10.1093/nar/gkh907) [Otto et al. (2014)](https://doi.org/10.1186/s12915-014-0086-0) | Antigenic |
| OG0003221 | Fam-b protein | It promotes the development of the parasite in both the liver and blood, either by supporting the development of the parasite within hepatocytes/erythrocytes and/or by manipulating the host's immune response. | [Fougère et al. (2016)](https://doi.org/10.1371/journal.ppat.1005917) | Discarded |
| OG0003438 | *Exported Plasmodium* proteins (8 PHISTa, 20 unknown) | PHIST family important for parasite-host interactions | [Yang et al. (2020](https://doi.org/10.3389/fmicb.2020.611190));  [Oberli et al. (2014)](https://doi.org/10.1096/fj.14-256057) | Antigenic |
| OG0003535 | PIR protein | Increased expression in stages that invade and reside in erythrocytes suggests an important role in parasite biology. | [Little et al. (2021](https://doi.org/10.1186/s12936-021-03979-6));  [Janssen et al. (2004);](https://doi.org/10.1093/nar/gkh907) [Otto et al. (2014)](https://doi.org/10.1186/s12915-014-0086-0) | Antigenic |
| OG0003627 | RIFIN | Antigen expressed on the surface of infected erythrocytes. Evidence suggests that it is important for evading the immune system. | [Bachmann et al. (2015);](https://doi.org/10.1186/s12936-015-0784-2)  [Saito et al. (2017);](https://doi.org/10.1038/nature24994)  [Goel et al. (2015)](https://doi.org/10.1038/nm.3812) | Antigenic |
| OG0003681 | PIR protein | Increased expression in stages that invade and reside in erythrocytes suggests an important role in parasite biology. | [Little et al. (2021](https://doi.org/10.1186/s12936-021-03979-6));  [Janssen et al. (2004);](https://doi.org/10.1093/nar/gkh907) [Otto et al. (2014)](https://doi.org/10.1186/s12915-014-0086-0) | Antigenic |
| OG0003734 | Exported protein (hyp11) | Linked to pathogenesis, antigenic variation and host cell remodeling. Its exact function is unknown. | [Pickford et al. (2021](https://hal.science/hal-03400342/document));  [Tachibana et al. (2012)](https://doi.org/10.1038/ng.2375) | Candidate |
| OG0003735 | PIR protein, YIR (1) | Increased expression in stages that invade and reside in erythrocytes suggests an important role in parasite biology. | [Little et al. (2021](https://doi.org/10.1186/s12936-021-03979-6));  [Janssen et al. (2004);](https://doi.org/10.1093/nar/gkh907) [Otto et al. (2014)](https://doi.org/10.1186/s12915-014-0086-0) | Antigenic |
| OG0003736 | PIR protein | Increased expression in stages that invade and reside in erythrocytes suggests an important role in parasite biology. | [Little et al. (2021](https://doi.org/10.1186/s12936-021-03979-6));  [Janssen et al. (2004);](https://doi.org/10.1093/nar/gkh907) [Otto et al. (2014)](https://doi.org/10.1186/s12915-014-0086-0) | Antigenic |
| **Gene Family** | **Product** | **Function** | **References** | **Classification** |
| OG0003795 | Fam-d protein | It is found in the erythrocyte membrane and partially colocalizes with the cytoplasmic domain of PfEMP1. It presents antigenic variation and generates immune response. | [Fernández-Becerra et al. (2020)](https://doi.org/10.1073/pnas.1920596117) | Antigenic |
| OG0003796 | PYST-C1 domain protein (fam-c) | Present in asexual stages that occur in red blood cells. | [Carlton et al. (2002)](https://doi.org/10.1038/nature01099) | Discarded |
| OG0003906 | STEVOR | It binds to red blood cells, promotes the invasion of merozoites or is present in stages where they invade and reside in erythrocytes. | [Niang et al. (2014](https://doi.org/10.1016/j.chom.2014.06.004)); [Bachmann et al. (2015)](https://doi.org/10.1186/s12936-015-0784-2) | Antigenic |
| OG0004062 | *Exported Plasmodium* protein | They can induce changes in the physiology of erythrocytes important for parasite growth and survival. | [Marti & Spielmann (2013)](https://doi.org/10.1016/j.mib.2013.04.010);  [Matthews et al. (2019)](https://doi.org/10.1111/cmi.13009) | Candidate |
| OG0004133 | PIR Protein (VIR) | Increased expression in stages that invade and reside in erythrocytes suggests an important role in parasite biology. | [Little et al. (2021](https://doi.org/10.1186/s12936-021-03979-6));  [Janssen et al. (2004);](https://doi.org/10.1093/nar/gkh907) [Otto et al. (2014)](https://doi.org/10.1186/s12915-014-0086-0) | Antigenic |
| OG0004137 | PIR protein | Increased expression in stages that invade and reside in erythrocytes suggests an important role in parasite biology. | [Little et al. (2021](https://doi.org/10.1186/s12936-021-03979-6));  [Janssen et al. (2004);](https://doi.org/10.1093/nar/gkh907) [Otto et al. (2014)](https://doi.org/10.1186/s12915-014-0086-0) | Antigenic |
| OG0004211 | SICAvar (type I) | Produced during the trophozoite stage and then inserted into the erythrocyte membrane and exposed to the outside of the host cell. | [Korir & Galinski (2006](https://doi.org/10.1016/j.meegid.2005.01.003));  [Howard & Barnwell (1984)](https://doi.org/10.1007/978-1-4684-4571-8_5) | Antigenic |
| OG0004292 | *Exported Plasmodium* protein (PHIST) | PHIST family important for parasite-host interactions | [Yang et al. (2020](https://doi.org/10.3389/fmicb.2020.611190)); [Oberli et al. (2014)](https://doi.org/10.1096/fj.14-256057) | Antigenic |
| OG0004293 | PIR Protein (CIR) | Increased expression in stages that invade and reside in erythrocytes suggests an important role in parasite biology. | [Little et al. (2021](https://doi.org/10.1186/s12936-021-03979-6));  [Janssen et al. (2004);](https://doi.org/10.1093/nar/gkh907) [Otto et al. (2014)](https://doi.org/10.1186/s12915-014-0086-0) | Antigenic |
| OG0004566 | *Exported Plasmodium* protein | They can induce changes in the physiology of erythrocytes important for parasite growth and survival. | [Marti & Spielmann (2013)](https://doi.org/10.1016/j.mib.2013.04.010);  [Matthews et al. (2019)](https://doi.org/10.1111/cmi.13009) | Candidate |
| OG0004567 | STP1 protein | They attach to red blood cells and intervene in the invasion process. They are related to the SURFIN family. | [Real et al. (2022)](https://doi.org/10.1016/j.mib.2022.102207) | Antigenic |
| OG0004656 | Pfmc-2TM protein | Present in infected erythrocytes during the trophozoite and schizont stage. Important for the transport to the surface of erythrocytes of proteins key to the virulence and survival of the parasite. | [Tsarukyanova et al. (2009](https://doi.org/10.1007/s00436-008-1270-3)); [Cooke et al. (2006)](https://doi.org/10.1083/jcb.200509122) | Antigenic |
| OG0004763 | PIR protein | Increased expression in stages that invade and reside in erythrocytes suggests an important role in parasite biology. | [Little et al. (2021](https://doi.org/10.1186/s12936-021-03979-6));  [Janssen et al. (2004);](https://doi.org/10.1093/nar/gkh907) [Otto et al. (2014)](https://doi.org/10.1186/s12915-014-0086-0) | Antigenic |
| **Gene Family** | **Product** | **Function** | **References** | **Classification** |
| OG0004868 | PIR protein | Increased expression in stages that invade and reside in erythrocytes suggests an important role in parasite biology. | [Little et al. (2021](https://doi.org/10.1186/s12936-021-03979-6));  [Janssen et al. (2004);](https://doi.org/10.1093/nar/gkh907) [Otto et al. (2014)](https://doi.org/10.1186/s12915-014-0086-0) | Antigenic |
| OG0004870 | RIFIN | Antigen expressed on the surface of infected erythrocytes. Evidence suggests that it is important for evading the immune system. | [Bachmann et al. (2015);](https://doi.org/10.1186/s12936-015-0784-2)  [Saito et al. (2017);](https://doi.org/10.1038/nature24994)  [Goel et al. (2015)](https://doi.org/10.1038/nm.3812) | Antigenic |
| OG0004962 | RIFIN | Antigen expressed on the surface of infected erythrocytes. Evidence suggests that it is important for evading the immune system. | [Bachmann et al. (2015);](https://doi.org/10.1186/s12936-015-0784-2)  [Saito et al. (2017);](https://doi.org/10.1038/nature24994)  [Goel et al. (2015)](https://doi.org/10.1038/nm.3812) | Antigenic |
| OG0005382 | PIR protein | Increased expression in stages that invade and reside in erythrocytes suggests an important role in parasite biology. | [Little et al. (2021](https://doi.org/10.1186/s12936-021-03979-6));  [Janssen et al. (2004);](https://doi.org/10.1093/nar/gkh907) [Otto et al. (2014)](https://doi.org/10.1186/s12915-014-0086-0) | Antigenic |
| OG0005513 | PIR protein | Increased expression in stages that invade and reside in erythrocytes suggests an important role in parasite biology. | [Little et al. (2021](https://doi.org/10.1186/s12936-021-03979-6));  [Janssen et al. (2004);](https://doi.org/10.1093/nar/gkh907) [Otto et al. (2014)](https://doi.org/10.1186/s12915-014-0086-0) | Antigenic |
| OG0005614 | Exported protein (PHISTb) | They are located in infected red blood cells and interact with the cytoskeleton of host cells. The RESA antigen is part of the PHISTb subfamily. | [Warncke et al. (2016); Sargent et al. (2006)](https://doi.org/10.1128/MMBR.00014-16) | Antigenic |
| OG0005616 | RIFIN | Antigen expressed on the surface of infected erythrocytes. Evidence suggests that it is important for evading the immune system. | [Bachmann et al. (2015);](https://doi.org/10.1186/s12936-015-0784-2)  [Saito et al. (2017);](https://doi.org/10.1038/nature24994)  [Goel et al. (2015)](https://doi.org/10.1038/nm.3812) | Antigenic |
| OG0005867 | PIR Protein (CIR) | Increased expression in stages that invade and reside in erythrocytes suggests an important role in parasite biology. | [Little et al. (2021](https://doi.org/10.1186/s12936-021-03979-6));  [Janssen et al. (2004);](https://doi.org/10.1093/nar/gkh907) [Otto et al. (2014)](https://doi.org/10.1186/s12915-014-0086-0) | Antigenic |
| OG0006027 | PIR protein | Increased expression in stages that invade and reside in erythrocytes suggests an important role in parasite biology. | [Little et al. (2021](https://doi.org/10.1186/s12936-021-03979-6));  [Janssen et al. (2004);](https://doi.org/10.1093/nar/gkh907) [Otto et al. (2014)](https://doi.org/10.1186/s12915-014-0086-0) | Antigenic |
| OG0006028 | PIR protein | Increased expression in stages that invade and reside in erythrocytes suggests an important role in parasite biology. | [Little et al. (2021](https://doi.org/10.1186/s12936-021-03979-6));  [Janssen et al. (2004);](https://doi.org/10.1093/nar/gkh907) [Otto et al. (2014)](https://doi.org/10.1186/s12915-014-0086-0) | Antigenic |
| OG0006029 | RIFIN | Antigen expressed on the surface of infected erythrocytes. Evidence suggests that it is important for evading the immune system. | [Bachmann et al. (2015);](https://doi.org/10.1186/s12936-015-0784-2)  [Saito et al. (2017);](https://doi.org/10.1038/nature24994)  [Goel et al. (2015)](https://doi.org/10.1038/nm.3812) | Antigenic |
| OG0006284 | RIFIN | Antigen expressed on the surface of infected erythrocytes. Evidence suggests that it is important for evading the immune system. | [Bachmann et al. (2015);](https://doi.org/10.1186/s12936-015-0784-2)  [Saito et al. (2017);](https://doi.org/10.1038/nature24994)  [Goel et al. (2015)](https://doi.org/10.1038/nm.3812) | Antigenic |
| OG0006582 | Chitinases | Necessary for the invasion of the midgut of the mosquito by mediating the penetration of the parasite through the peritrophic membrane. | [Viswanath et al. (2021)](https://doi.org/10.1002/pro.4095) | Antigenic |
| OG0006954 | PIR protein | Increased expression in stages that invade and reside in erythrocytes suggests an important role in parasite biology. | [Little et al. (2021](https://doi.org/10.1186/s12936-021-03979-6));  [Janssen et al. (2004);](https://doi.org/10.1093/nar/gkh907) [Otto et al. (2014)](https://doi.org/10.1186/s12915-014-0086-0) | Antigenic |
| OG0006955 | RIFIN | Antigen expressed on the surface of infected erythrocytes. Evidence suggests that it is important for evading the immune system. | [Bachmann et al. (2015);](https://doi.org/10.1186/s12936-015-0784-2)  [Saito et al. (2017);](https://doi.org/10.1038/nature24994)  [Goel et al. (2015)](https://doi.org/10.1038/nm.3812) | Antigenic |
| **Gene Family** | **Product** | **Function** | **References** | **Classification** |
| OG0006956 | PIR protein | Increased expression in stages that invade and reside in erythrocytes suggests an important role in parasite biology. | [Little et al. (2021](https://doi.org/10.1186/s12936-021-03979-6));  [Janssen et al. (2004);](https://doi.org/10.1093/nar/gkh907) [Otto et al. (2014)](https://doi.org/10.1186/s12915-014-0086-0) | Antigenic |
| OG0007772 | *Plasmodium* conserved membrane protein | Some keys to the maturation of exo-erythrocytic forms, others are not expressed in stages that occur in red blood cells. | [Al-Nihmi et al. (2017); Kaiser et al. (2004)](https://doi.org/10.1038/srep40407) | Discarded |
| OG0007776 | STEVOR | It binds to red blood cells, promotes the invasion of merozoites or is present in stages where they invade and reside in erythrocytes. | [Niang et al. (2014](https://doi.org/10.1016/j.chom.2014.06.004)); [Bachmann et al. (2015)](https://doi.org/10.1186/s12936-015-0784-2) | Antigenic |
| OG0007777 | PIR protein | Increased expression in stages that invade and reside in erythrocytes suggests an important role in parasite biology. | [Little et al. (2021](https://doi.org/10.1186/s12936-021-03979-6));  [Janssen et al. (2004);](https://doi.org/10.1093/nar/gkh907) [Otto et al. (2014)](https://doi.org/10.1186/s12915-014-0086-0) | Antigenic |
| OG0008551 | Agamete ntígeno 27/25 | High expression after invasion by merozoites during gametocyte synthesis. During the infection process, it is an immunogenic antigen. | [Carter et al. (1989)](https://doi.org/10.1016/0014-4894(89)90182-3) | Antigenic |
| OG0008558 | RIFIN | Antigen expressed on the surface of infected erythrocytes. Evidence suggests that it is important for evading the immune system. | [Bachmann et al. (2015);](https://doi.org/10.1186/s12936-015-0784-2)  [Saito et al. (2017);](https://doi.org/10.1038/nature24994)  [Goel et al. (2015)](https://doi.org/10.1038/nm.3812) | Antigenic |
| OG0008764 | KELT protein | Telomere gene in P. *falciparum.* Expanded in tandem matrix in *P. ovale.* Signal peptide in versions of *P. ovale.* | [Ansari et al. (2016)](https://doi.org/10.1016/j.ijpara.2016.05.009) | Candidate |
| OG0008942 | EPF3 | They are exported to the Maurer cleft, suggesting a possible role in aiding the correct presentation of membrane proteins on the surface of infected erythrocytes. | [Liu et al. (2018); Mbengue et al. (2013)](https://doi.org/10.1186/s12864-018-4654-5) | Antigenic |
| OG0009146 | Fam-a protein | Important for the invasion of erythrocytes, expressed in merozoites, keys to the development of the parasite. | [Alam et al. (2016](https://doi.org/10.1016/j.bbrc.2016.08.096));  [Zeeshan et al. (2015)](https://doi.org/10.1093/infdis/jiu558) | Antigenic |
| OG0009148 | Protein exported from *Plasmodium*, hypothetical protein | They can induce changes in the physiology of erythrocytes important for parasite growth and survival. | [Marti & Spielmann (2013)](https://doi.org/10.1016/j.mib.2013.04.010);  [Matthews et al. (2019)](https://doi.org/10.1111/cmi.13009) | Candidate |
| OG0009152 | PIR protein | Increased expression in stages that invade and reside in erythrocytes suggests an important role in parasite biology. | [Little et al. (2021](https://doi.org/10.1186/s12936-021-03979-6));  [Janssen et al. (2004);](https://doi.org/10.1093/nar/gkh907) [Otto et al. (2014)](https://doi.org/10.1186/s12915-014-0086-0) | Antigenic |
| OG0009359 | STEVOR | It binds to red blood cells, promotes the invasion of merozoites or is present in stages where they invade and reside in erythrocytes. | [Niang et al. (2014](https://doi.org/10.1016/j.chom.2014.06.004)); [Bachmann et al. (2015)](https://doi.org/10.1186/s12936-015-0784-2) | Antigenic |
| OG0009360 | PfEMP1 (28), non-specific product (1) | It undergoes antigenic variation to evade the immune system and changes the cytoadhesive properties of infected red blood cells so that they bind to uninfected ones. | [Pasternak & Dzikowski (2009](https://doi.org/10.1016/j.biocel.2008.12.012));  [Jensen et al. (2019)](https://doi.org/10.1111/imr.12807) | Antigenic |
| **Gene Family** | **Product** | **Function** | **References** | **Classification** |
| OG0009361 | Fam-b protein (15), non-specific product (2) | It promotes the development of the parasite in both the liver and blood, either by supporting the development of the parasite within hepatocytes/erythrocytes and/or by manipulating the host's immune response. | [Fougère et al. (2016)](https://doi.org/10.1371/journal.ppat.1005917) | Discarded |
| OG0009362 | Fam-a protein (24), non-specific product (1) | Important for the invasion of erythrocytes, expressed in merozoites, keys to the development of the parasite. | [Alam et al. (2016](https://doi.org/10.1016/j.bbrc.2016.08.096));  [Zeeshan et al. (2015)](https://doi.org/10.1093/infdis/jiu558) | Antigenic |
| OG0009364 | PIR protein | Increased expression in stages that invade and reside in erythrocytes suggests an important role in parasite biology. | [Little et al. (2021](https://doi.org/10.1186/s12936-021-03979-6));  [Janssen et al. (2004);](https://doi.org/10.1093/nar/gkh907) [Otto et al. (2014)](https://doi.org/10.1186/s12915-014-0086-0) | Antigenic |
| OG0009580 | PfEMP1 | It undergoes antigenic variation to evade the immune system and changes the cytoadhesive properties of infected red blood cells so that they bind to uninfected ones. | [Pasternak & Dzikowski (2009](https://doi.org/10.1016/j.biocel.2008.12.012));  [Jensen et al. (2019)](https://doi.org/10.1111/imr.12807) | Antigenic |
| OG0009583 | PIR protein (KIR, type VIR7) | Increased expression in stages that invade and reside in erythrocytes suggests an important role in parasite biology. | [Little et al. (2021](https://doi.org/10.1186/s12936-021-03979-6));  [Janssen et al. (2004);](https://doi.org/10.1093/nar/gkh907) [Otto et al. (2014)](https://doi.org/10.1186/s12915-014-0086-0) | Antigenic |
| OG0009585 | EMP1 | It undergoes antigenic variation to evade the immune system and changes the cytoadhesive properties of infected red blood cells so that they bind to uninfected ones. | [Pasternak & Dzikowski (2009](https://doi.org/10.1016/j.biocel.2008.12.012));  [Jensen et al. (2019)](https://doi.org/10.1111/imr.12807) | Antigenic |
| OG0009851 | STEVOR | It binds to red blood cells, promotes the invasion of merozoites or is present in stages where they invade and reside in erythrocytes. | [Niang et al. (2014](https://doi.org/10.1016/j.chom.2014.06.004)); [Bachmann et al. (2015)](https://doi.org/10.1186/s12936-015-0784-2) | Antigenic |
| OG0010136 | PIR protein | Increased expression in stages that invade and reside in erythrocytes suggests an important role in parasite biology. | [Little et al. (2021](https://doi.org/10.1186/s12936-021-03979-6));  [Janssen et al. (2004);](https://doi.org/10.1093/nar/gkh907) [Otto et al. (2014)](https://doi.org/10.1186/s12915-014-0086-0) | Antigenic |
| OG0010444 | *Exported Plasmodium* protein | They can induce changes in the physiology of erythrocytes important for parasite growth and survival. | [Marti & Spielmann (2013)](https://doi.org/10.1016/j.mib.2013.04.010);  [Matthews et al. (2019)](https://doi.org/10.1111/cmi.13009) | Candidate |
| OG0010453 | PIR protein | Increased expression in stages that invade and reside in erythrocytes suggests an important role in parasite biology. | [Little et al. (2021](https://doi.org/10.1186/s12936-021-03979-6));  [Janssen et al. (2004);](https://doi.org/10.1093/nar/gkh907) [Otto et al. (2014)](https://doi.org/10.1186/s12915-014-0086-0) | Antigenic |
| OG0010454 | Exported Plasmodium protein (PHIST) | PHIST family important for parasite-host interactions | [Yang et al. (2020](https://doi.org/10.3389/fmicb.2020.611190));  [Oberli et al. (2014)](https://doi.org/10.1096/fj.14-256057) | Antigenic |
| OG0010455 | PIR protein | Increased expression in stages that invade and reside in erythrocytes suggests an important role in parasite biology. | [Little et al. (2021](https://doi.org/10.1186/s12936-021-03979-6));  [Janssen et al. (2004);](https://doi.org/10.1093/nar/gkh907) [Otto et al. (2014)](https://doi.org/10.1186/s12915-014-0086-0) | Antigenic |
| OG0010458 | *Exported Plasmodium* protein | They can induce changes in the physiology of erythrocytes important for parasite growth and survival. | [Marti & Spielmann (2013)](https://doi.org/10.1016/j.mib.2013.04.010);  [Matthews et al. (2019)](https://doi.org/10.1111/cmi.13009) | Candidate |
| **Gene Family** | **Product** | **Function** | **References** | **Classification** |
| OG0010750 | Protein exported from *Plasmodium*, hypothetical protein | They can induce changes in the physiology of erythrocytes important for parasite growth and survival. | [Marti & Spielmann (2013)](https://doi.org/10.1016/j.mib.2013.04.010);  [Matthews et al. (2019)](https://doi.org/10.1111/cmi.13009) | Candidate |
| OG0010751 | PIR protein | Increased expression in stages that invade and reside in erythrocytes suggests an important role in parasite biology. | [Little et al. (2021](https://doi.org/10.1186/s12936-021-03979-6));  [Janssen et al. (2004);](https://doi.org/10.1093/nar/gkh907) [Otto et al. (2014)](https://doi.org/10.1186/s12915-014-0086-0) | Antigenic |
| OG0011086 | RIFIN | Antigen expressed on the surface of infected erythrocytes. Evidence suggests that it is important for evading the immune system. | [Bachmann et al. (2015);](https://doi.org/10.1186/s12936-015-0784-2)  [Saito et al. (2017);](https://doi.org/10.1038/nature24994)  [Goel et al. (2015)](https://doi.org/10.1038/nm.3812) | Antigenic |
| OG0011088 | PIR protein (KIR, type VIR7) | Increased expression in stages that invade and reside in erythrocytes suggests an important role in parasite biology. | [Little et al. (2021](https://doi.org/10.1186/s12936-021-03979-6));  [Janssen et al. (2004);](https://doi.org/10.1093/nar/gkh907) [Otto et al. (2014)](https://doi.org/10.1186/s12915-014-0086-0) | Antigenic |
| OG0011092 | PIR protein | Increased expression in stages that invade and reside in erythrocytes suggests an important role in parasite biology. | [Little et al. (2021](https://doi.org/10.1186/s12936-021-03979-6));  [Janssen et al. (2004);](https://doi.org/10.1093/nar/gkh907) [Otto et al. (2014)](https://doi.org/10.1186/s12915-014-0086-0) | Antigenic |
| OG0011451 | *Exported Plasmodium* protein (PHISTa) | PHIST family important for parasite-host interactions | [Yang et al. (2020](https://doi.org/10.3389/fmicb.2020.611190)); [Oberli et al. (2014)](https://doi.org/10.1096/fj.14-256057) | Antigenic |
| OG0011460 | PfEMP1/EMP1, cytoadhesion-linked asexual protein 3.1 (1) | Functions in cytoadhesion processes key to the invasion process and therefore, the virulence of the parasite. | [Pasternak & Dzikowski (2009](https://doi.org/10.1016/j.biocel.2008.12.012));  [Jensen et al. (2019);](https://doi.org/10.1111/imr.12807) [Ocampo et al. (2005)](https://doi.org/10.1110/ps.04883905) | Antigenic |
| OG0011463 | PIR protein | Increased expression in stages that invade and reside in erythrocytes suggests an important role in parasite biology. | [Little et al. (2021](https://doi.org/10.1186/s12936-021-03979-6));  [Janssen et al. (2004);](https://doi.org/10.1093/nar/gkh907) [Otto et al. (2014)](https://doi.org/10.1186/s12915-014-0086-0) | Antigenic |
| OG0011464 | PIR Protein (VIR) | Increased expression in stages that invade and reside in erythrocytes suggests an important role in parasite biology. | [Little et al. (2021](https://doi.org/10.1186/s12936-021-03979-6));  [Janssen et al. (2004);](https://doi.org/10.1093/nar/gkh907) [Otto et al. (2014)](https://doi.org/10.1186/s12915-014-0086-0) | Antigenic |
| OG0012882 | PIR protein | Increased expression in stages that invade and reside in erythrocytes suggests an important role in parasite biology. | [Little et al. (2021](https://doi.org/10.1186/s12936-021-03979-6));  [Janssen et al. (2004);](https://doi.org/10.1093/nar/gkh907) [Otto et al. (2014)](https://doi.org/10.1186/s12915-014-0086-0) | Antigenic |
| OG0012884 | PIR protein (KIR, CYIR, KIR type, Vir7 type) | Increased expression in stages that invade and reside in erythrocytes suggests an important role in parasite biology. | [Little et al. (2021](https://doi.org/10.1186/s12936-021-03979-6));  [Janssen et al. (2004);](https://doi.org/10.1093/nar/gkh907) [Otto et al. (2014)](https://doi.org/10.1186/s12915-014-0086-0) | Antigenic |
| OG0012886 | *Exported Plasmodium* protein | They can induce changes in the physiology of erythrocytes important for parasite growth and survival. | [Marti & Spielmann (2013)](https://doi.org/10.1016/j.mib.2013.04.010);  [Matthews et al. (2019)](https://doi.org/10.1111/cmi.13009) | Candidate |
| OG0012887 | Fam-l protein | They appear to be exported from the parasite to blood cells. | [Rutledge et al. (2017)](https://www.nature.com/articles/nature21038) | Antigenic |
| OG0013479 | RIFIN | Antigen expressed on the surface of infected erythrocytes. Evidence suggests that it is important for evading the immune system. | [Bachmann et al. (2015);](https://doi.org/10.1186/s12936-015-0784-2)  [Saito et al. (2017);](https://doi.org/10.1038/nature24994)  [Goel et al. (2015)](https://doi.org/10.1038/nm.3812) | Antigenic |
| **Gene Family** | **Product** | **Function** | **References** | **Classification** |
| OG0013489 | PIR Protein (CIR) | Increased expression in stages that invade and reside in erythrocytes suggests an important role in parasite biology. | [Little et al. (2021](https://doi.org/10.1186/s12936-021-03979-6));  [Janssen et al. (2004);](https://doi.org/10.1093/nar/gkh907) [Otto et al. (2014)](https://doi.org/10.1186/s12915-014-0086-0) | Antigenic |
| OG0014073 | *Exported Plasmodium* protein (PHISTa) | PHIST family important for parasite-host interactions, expressed in infected red blood cells. | [Yang et al. (2020](https://doi.org/10.3389/fmicb.2020.611190)); [Oberli et al. (2014)](https://doi.org/10.1096/fj.14-256057) | Antigenic |
| OG0014080 | PIR protein | Increased expression in stages that invade and reside in erythrocytes suggests an important role in parasite biology. | [Little et al. (2021](https://doi.org/10.1186/s12936-021-03979-6));  [Janssen et al. (2004);](https://doi.org/10.1093/nar/gkh907) [Otto et al. (2014)](https://doi.org/10.1186/s12915-014-0086-0) | Antigenic |
| OG0014081 | PIR protein | Increased expression in stages that invade and reside in erythrocytes suggests an important role in parasite biology. | [Little et al. (2021](https://doi.org/10.1186/s12936-021-03979-6));  [Janssen et al. (2004);](https://doi.org/10.1093/nar/gkh907) [Otto et al. (2014)](https://doi.org/10.1186/s12915-014-0086-0) | Antigenic |
| OG0015533 | RESA N-terminal, RAD (Pv-fam-e), non-specific product (1) | Important for the survival of the parasite and the process of infection. RAD with structure similar to PHIST proteins. | [Pei et al. (2007](https://doi.org/10.1182/blood-2007-02-076919));  [Rug & Maier (2011); Carlton et al. (2008);](https://doi.org/10.1002/iub.525)  [Warncke et al. (2016)](https://doi.org/10.1128/MMBR.00014-16) | Antigenic |
| OG0015534 | SICAvar (type I, II) | Produced during the trophozoite stage and then inserted into the erythrocyte membrane and exposed to the outside of the host cell. | [Korir & Galinski (2006](https://doi.org/10.1016/j.meegid.2005.01.003));  [Howard & Barnwell (1984)](https://doi.org/10.1007/978-1-4684-4571-8_5) | Antigenic |
| OG0015535 | SICAvar (type II) | Produced during the trophozoite stage and then inserted into the erythrocyte membrane and exposed to the outside of the host cell. | [Korir & Galinski (2006](https://doi.org/10.1016/j.meegid.2005.01.003));  [Howard & Barnwell (1984)](https://doi.org/10.1007/978-1-4684-4571-8_5) | Antigenic |
| OG0015537 | PIR protein | Increased expression in stages that invade and reside in erythrocytes suggests an important role in parasite biology. | [Little et al. (2021](https://doi.org/10.1186/s12936-021-03979-6));  [Janssen et al. (2004);](https://doi.org/10.1093/nar/gkh907) [Otto et al. (2014)](https://doi.org/10.1186/s12915-014-0086-0) | Antigenic |
| OG0016368 | *Exported Plasmodium* protein | They can induce changes in the physiology of erythrocytes important for parasite growth and survival. | [Marti & Spielmann (2013)](https://doi.org/10.1016/j.mib.2013.04.010);  [Matthews et al. (2019)](https://doi.org/10.1111/cmi.13009) | Candidate |
| OG0016383 | Ubiquitin carboxyl-terminal hydrolase, PIR (KIR, type-KIR) | UCH is likely essential for parasite growth and survival. PIR proteins present in asexual stages that occur in erythrocytes. | [Artavanis-Tsakonas et al. (2010);](https://doi.org/10.1074/jbc.M109.072405)  [Little et al. (2021);](https://doi.org/10.1186/s12936-021-03979-6)  [Janssen et al. (2004);](https://doi.org/10.1093/nar/gkh907) [Otto et al. (2014)](https://doi.org/10.1186/s12915-014-0086-0) | Antigenic |
| OG0016398 | *Exported Plasmodium* protein | They can induce changes in the physiology of erythrocytes important for parasite growth and survival. | [Marti & Spielmann (2013)](https://doi.org/10.1016/j.mib.2013.04.010);  [Matthews et al. (2019)](https://doi.org/10.1111/cmi.13009) | Candidate |
| OG0016406 | PIR Protein (CIR) | Increased expression in stages that invade and reside in erythrocytes suggests an important role in parasite biology. | [Little et al. (2021](https://doi.org/10.1186/s12936-021-03979-6));  [Janssen et al. (2004);](https://doi.org/10.1093/nar/gkh907) [Otto et al. (2014)](https://doi.org/10.1186/s12915-014-0086-0) | Antigenic |
| OG0017378 | Fam-b protein | It promotes the development of the parasite in both the liver and blood, either by supporting the development of the parasite within hepatocytes/erythrocytes and/or by manipulating the host's immune response. | [Fougère et al. (2016)](https://doi.org/10.1371/journal.ppat.1005917) | Discarded |
| OG0017399 | Hypothetical protein | - | - | Discarded |
| **Gene Family** | **Product** | **Function** | **References** | **Classification** |
| OG0018679 | Fam-a protein | Important for the invasion of erythrocytes, expressed in merozoites, keys to the development of the parasite. | [Alam et al. (2016](https://doi.org/10.1016/j.bbrc.2016.08.096));  [Zeeshan et al. (2015)](https://doi.org/10.1093/infdis/jiu558) | Antigenic |
| OG0020391 | MSP3 | Expressed secreted merozoite surface protein (antigen) important for the invasion of erythrocytes. | [Deshmukh et al. (2018)](https://doi.org/10.1128/IAI.00067-18) | Antigenic |
| OG0022865 | PIR protein | Increased expression in stages that invade and reside in erythrocytes suggests an important role in parasite biology. | [Little et al. (2021](https://doi.org/10.1186/s12936-021-03979-6));  [Janssen et al. (2004);](https://doi.org/10.1093/nar/gkh907) [Otto et al. (2014)](https://doi.org/10.1186/s12915-014-0086-0) | Antigenic |
| OG0030154 | Hypothetical protein | - | - | Discarded |
| OG0030213 | PfEMP1 (5), hypothetical protein (1) | It undergoes antigenic variation to evade the immune system and changes the cytoadhesive properties of infected red blood cells so that they bind to uninfected ones. | [Pasternak & Dzikowski (2009](https://doi.org/10.1016/j.biocel.2008.12.012));  [Jensen et al. (2019)](https://doi.org/10.1111/imr.12807) | Antigenic |
| OG0034644 | PIR protein | Increased expression in stages that invade and reside in erythrocytes suggests an important role in parasite biology. | [Little et al. (2021](https://doi.org/10.1186/s12936-021-03979-6));  [Janssen et al. (2004);](https://doi.org/10.1093/nar/gkh907) [Otto et al. (2014)](https://doi.org/10.1186/s12915-014-0086-0) | Antigenic |
| OG0040450 | Hypothetical protein | - | - | Discarded |
