## Supplementary material for "Tracking the intensity of the mechanism to produce antigenic diversity by subtelomeric ectopic recombination across the phylogeny of *Plasmodium* parasites": Supplementary_material_dataset6.pdf

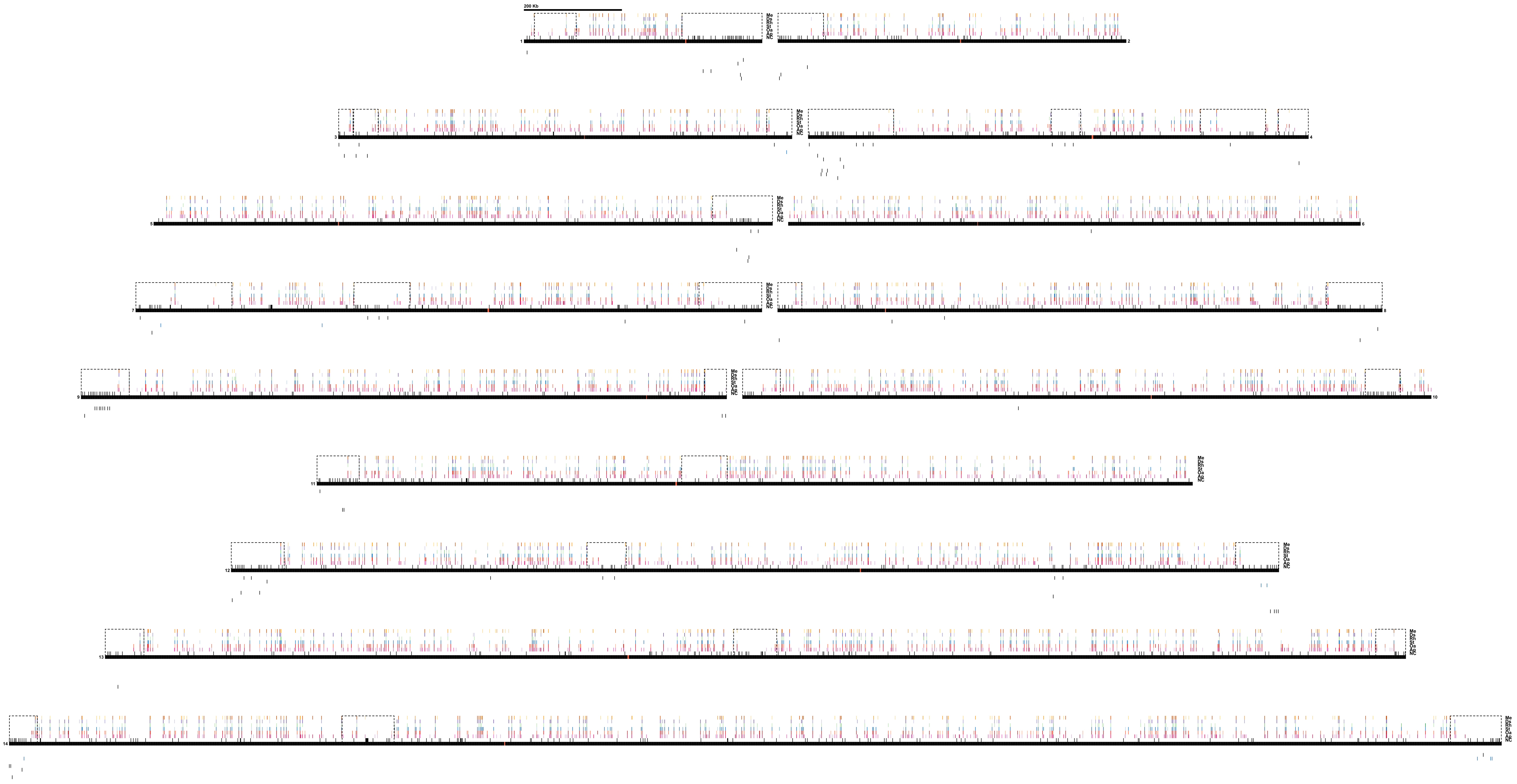

200 Kb
Me
Ds
Rh
St
Oa
Ap
NC
1
2
3
4
5
6
7
8
9
10
11
12
13
14

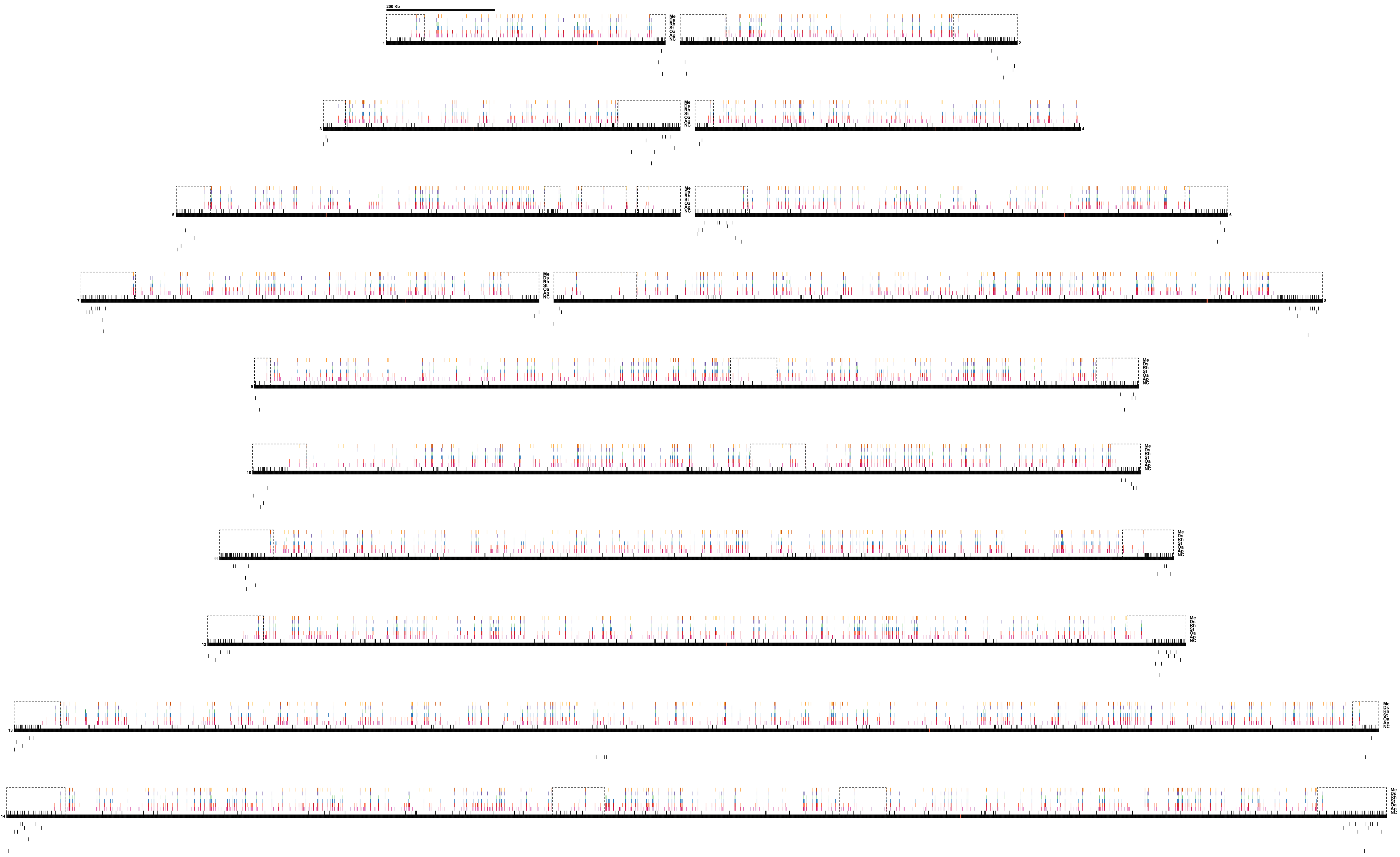
200 Kb
Me
Ds
Rh
St
Oa
Ap
NC
1
2
3
4
5
6
7
8
9
10
11
12
13
14

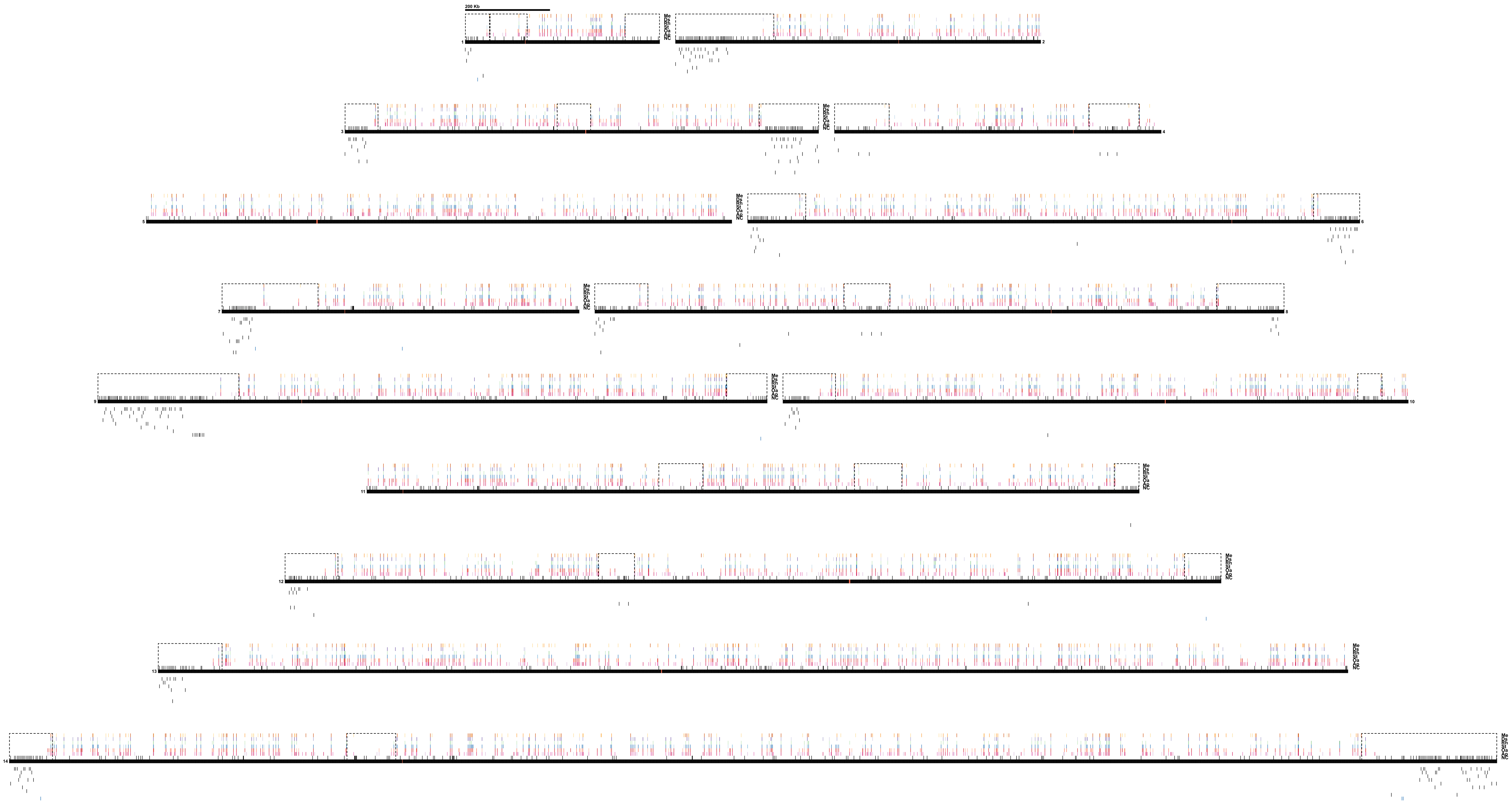
200 Kb
Me
Ds
Rh
St
Oa
Ap
NC
1
2
3
4
5
6
7
8
9
10
11
12
13
14

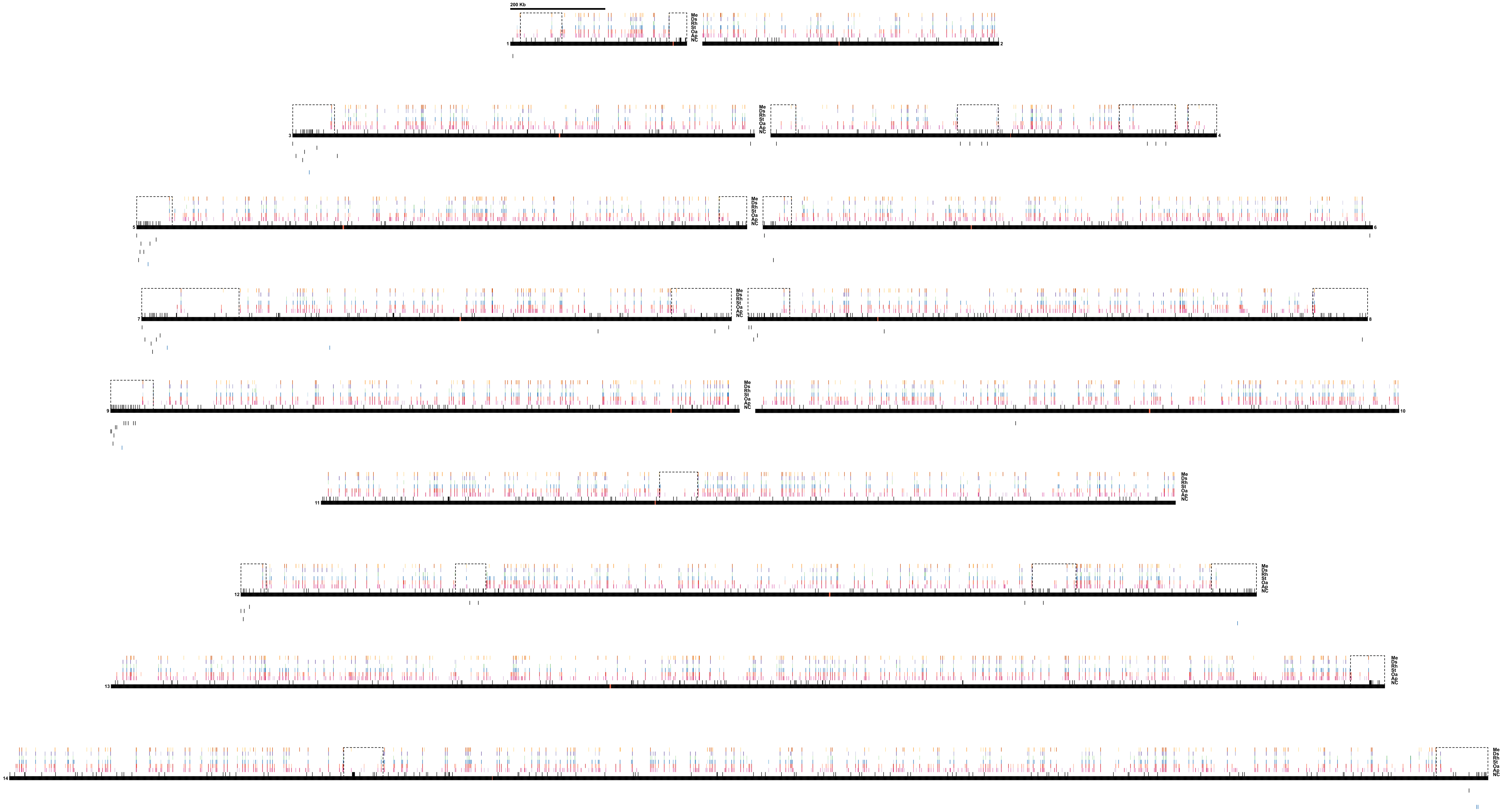
200 Kb
Me
Ds
Rh
St
Oa
Ap
NC
1
2
3
4
5
6
7
8
9
10
11
12
13
14

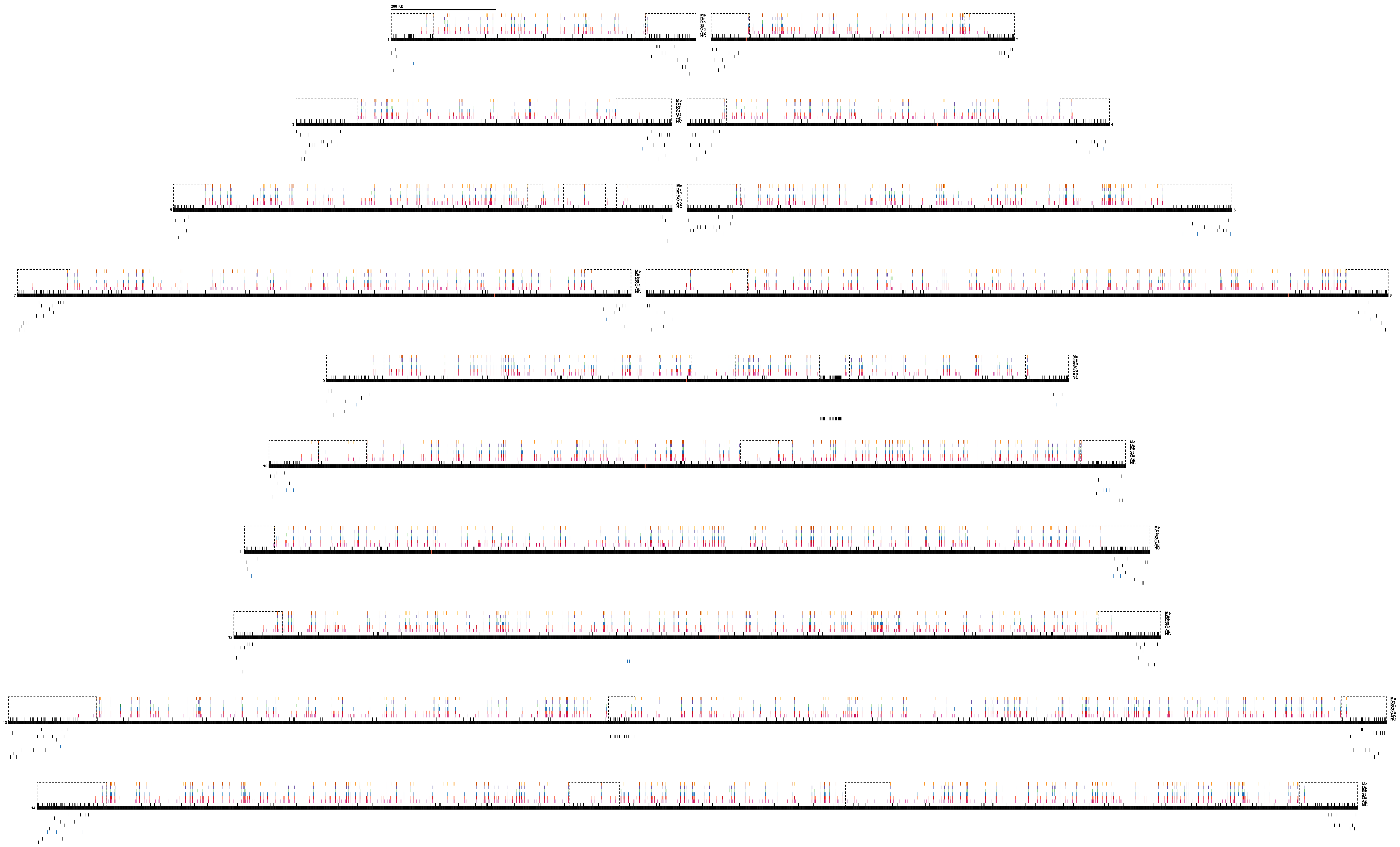
200 Kb
Me
Ds
Rh
St
Oa
Ap
NC
1
2
3
4
5
6
7
8
9
10
11
12
13
14

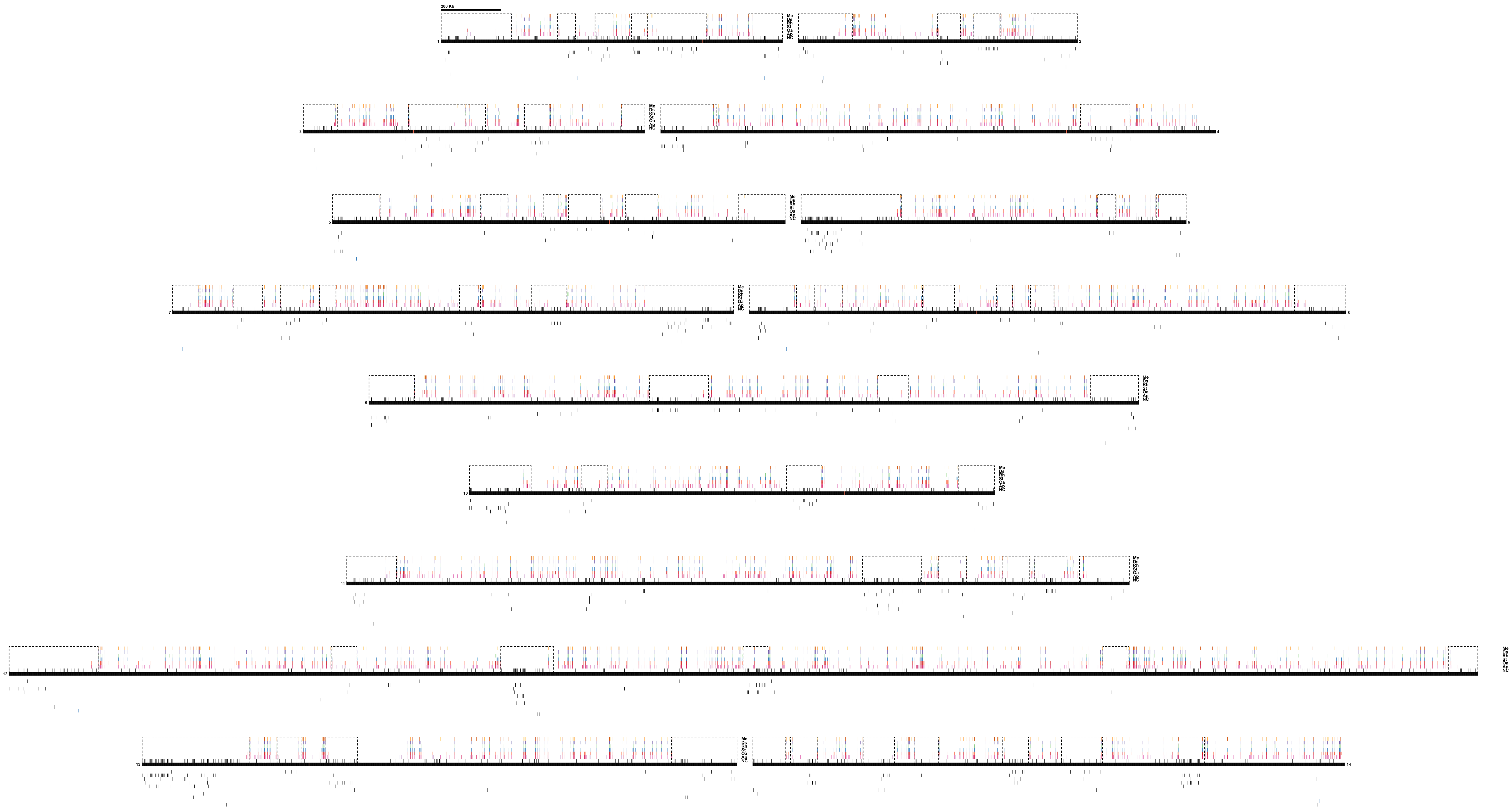
200 Kb
Me
Ds
Rh
St
Oa
Ap
NC
1
2
3
4
5
6
7
8
9
10
11
12
13
14

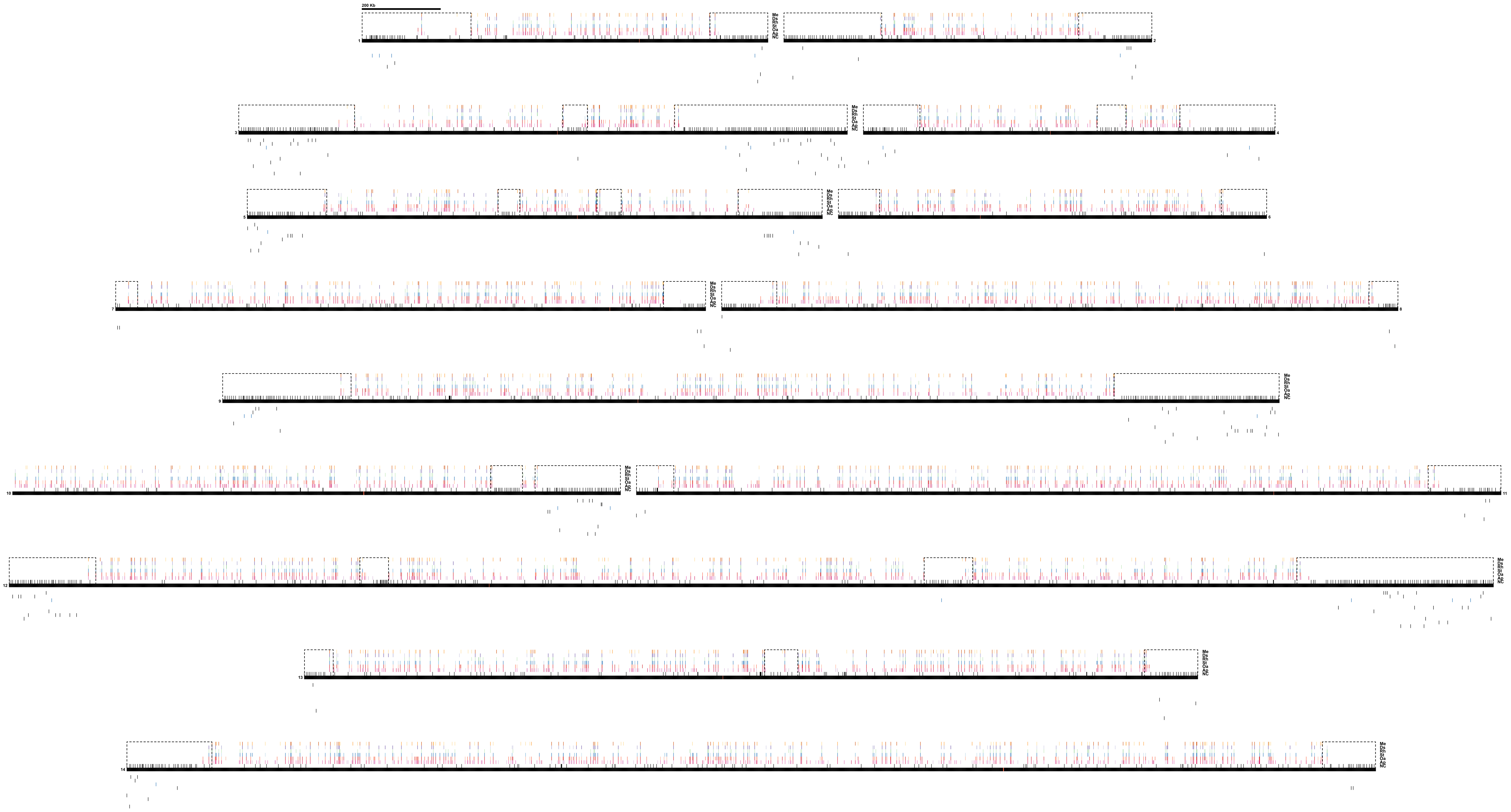
200 Kb

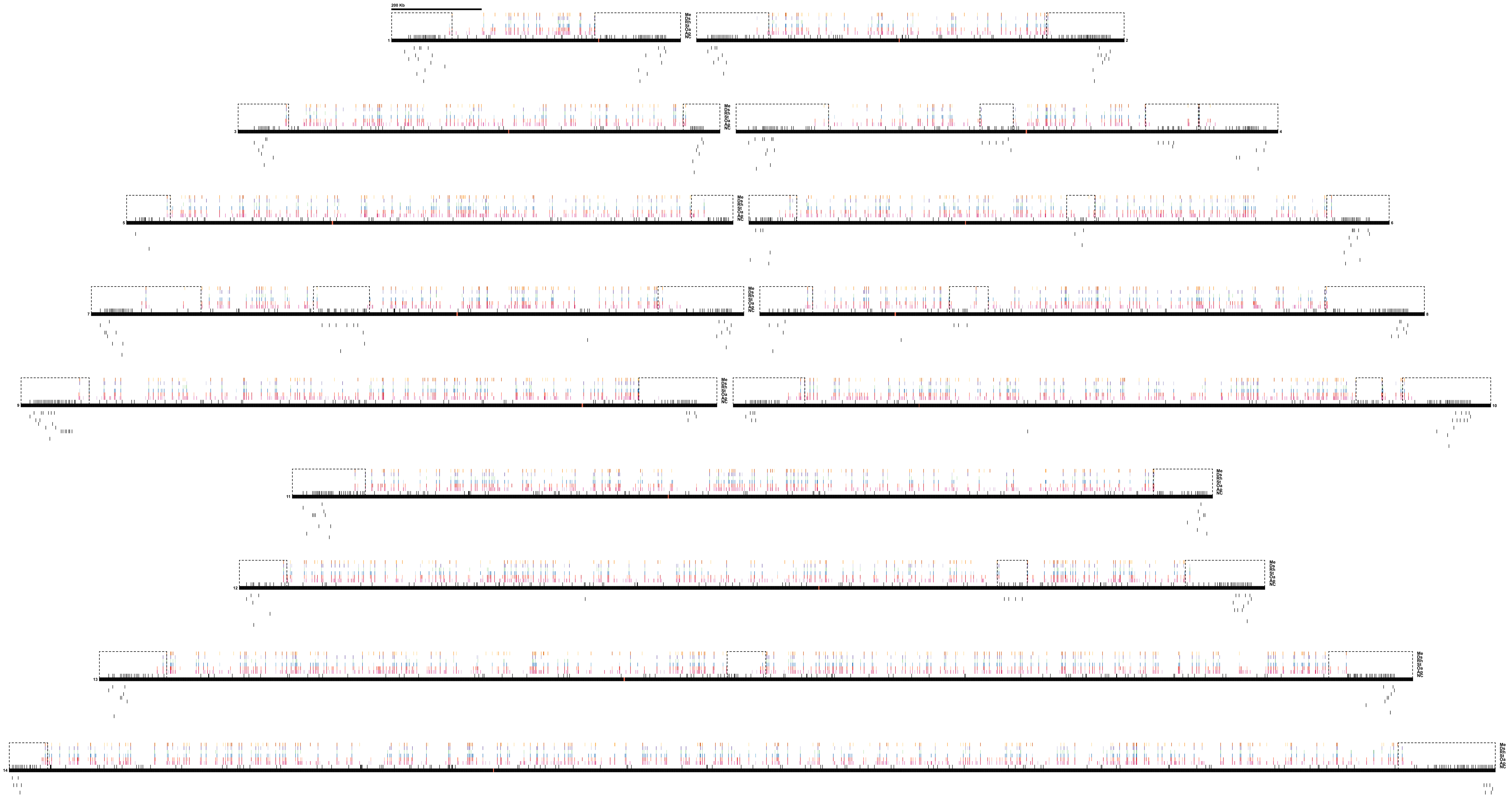
200 Kb
Me
Ds
Rh
St
Oa
Ap
NC
1
2
3
4
5
6
7
8
9
10
11
12
13
14

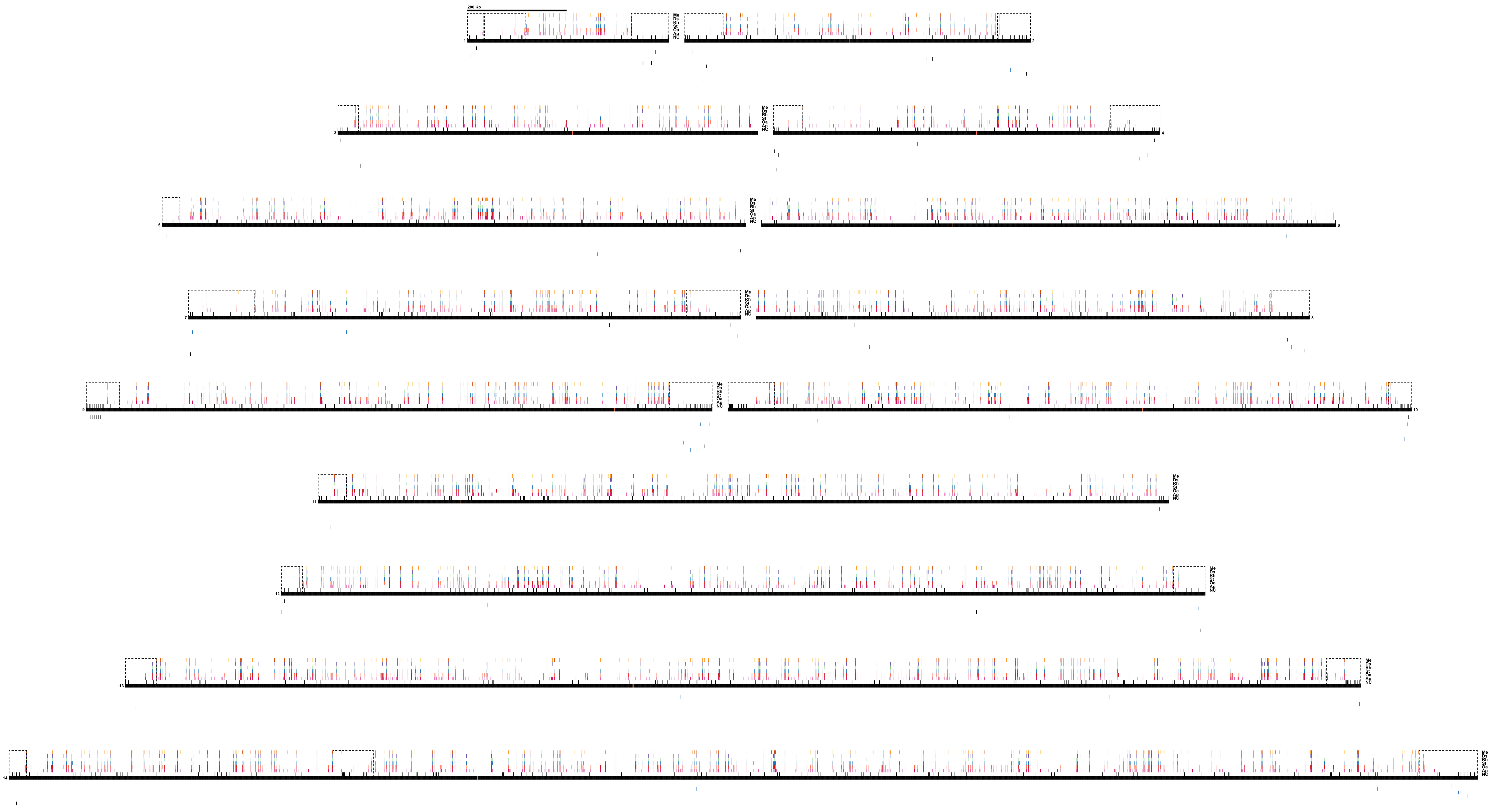

200 Kb
Me
Ds
Rh
St
Oa
Ap
NC
1
2
3
4
5
6
7
8
9
10
11
12
13
14

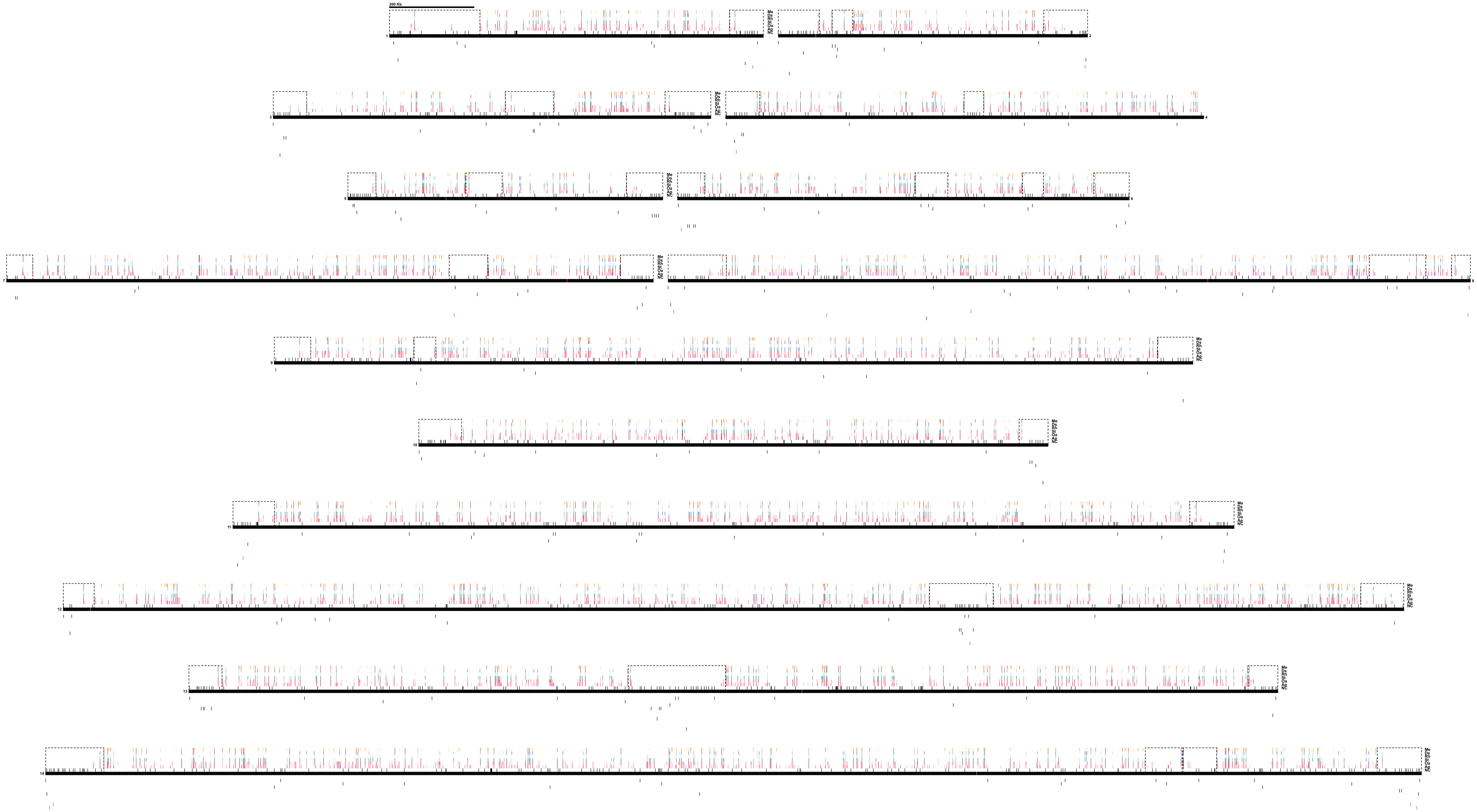

200 Kb
Me
Ds
Rh
St
Oa
Ap
NC
1
2
3
4
5
6
7
8
9
10
11
12
13
14

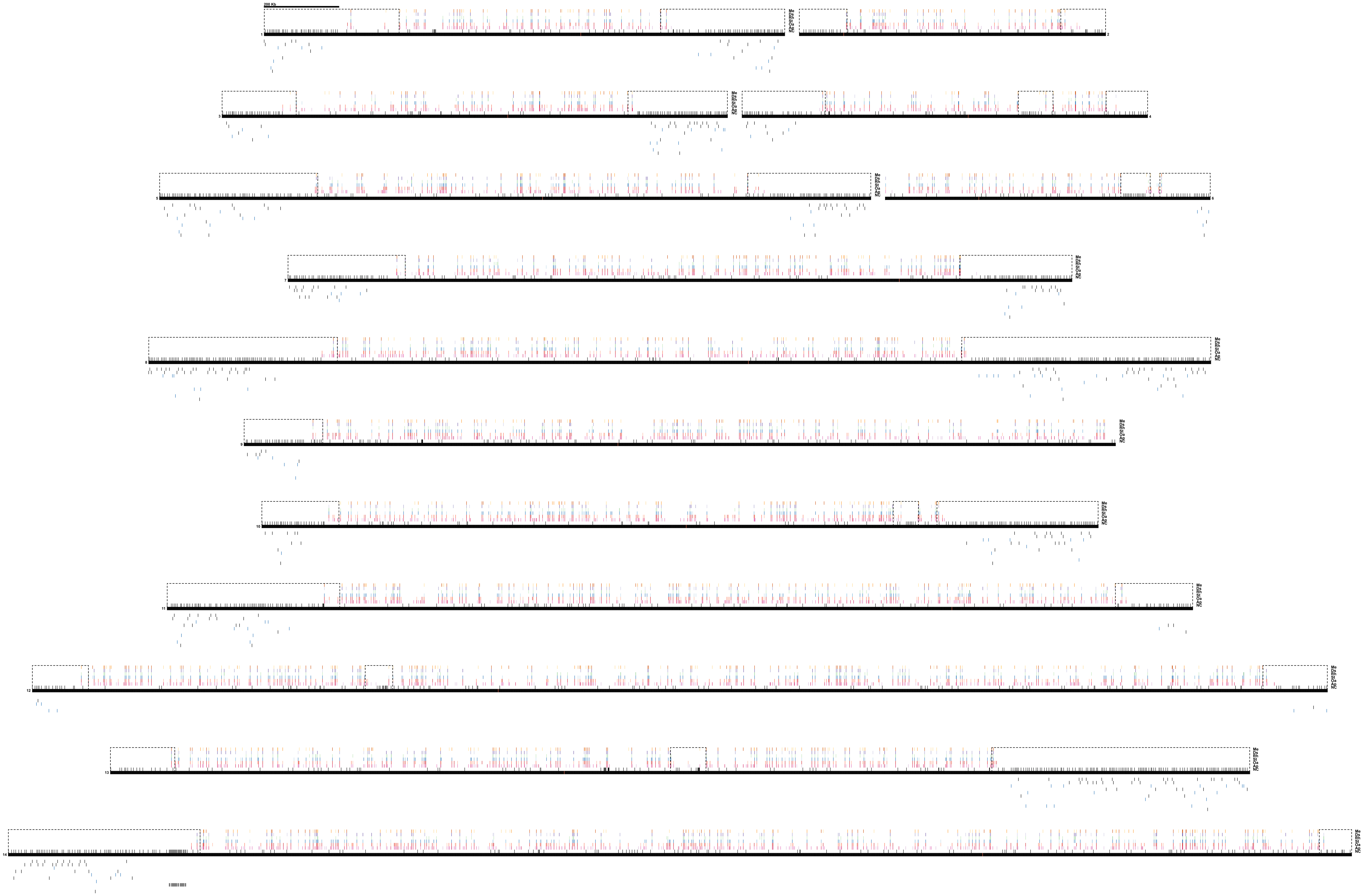
200 Kb
Me
Ds
Rh
St
Oa
Ap
NC
1
2
3
4
5
6
7
8
9
10
11
12
13
14

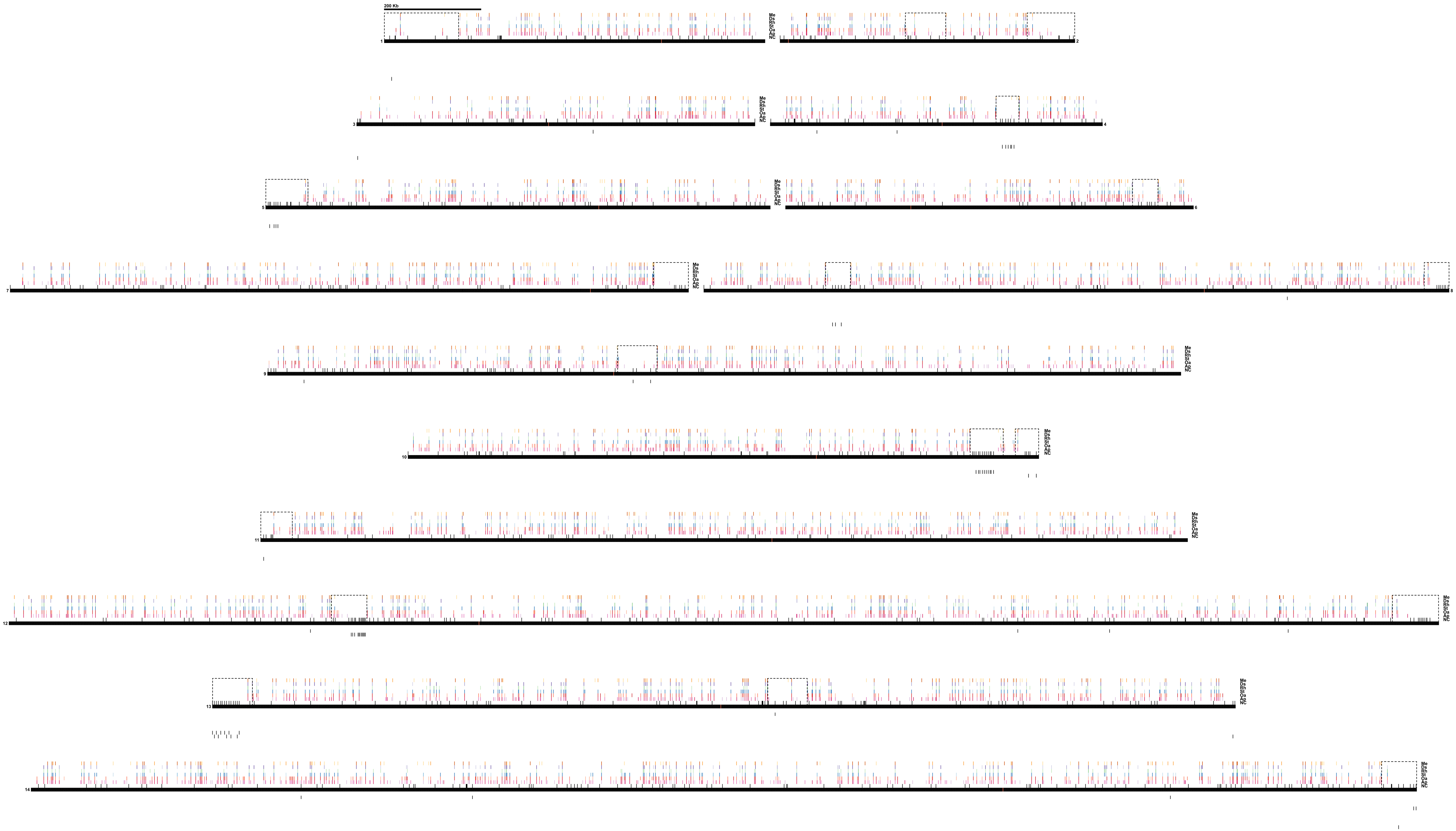
200 Kb
Me
Ds
Rh
St
Oa
Ap
NC
1
2
3
4
5
6
7
8
9
10
11
12
13
14

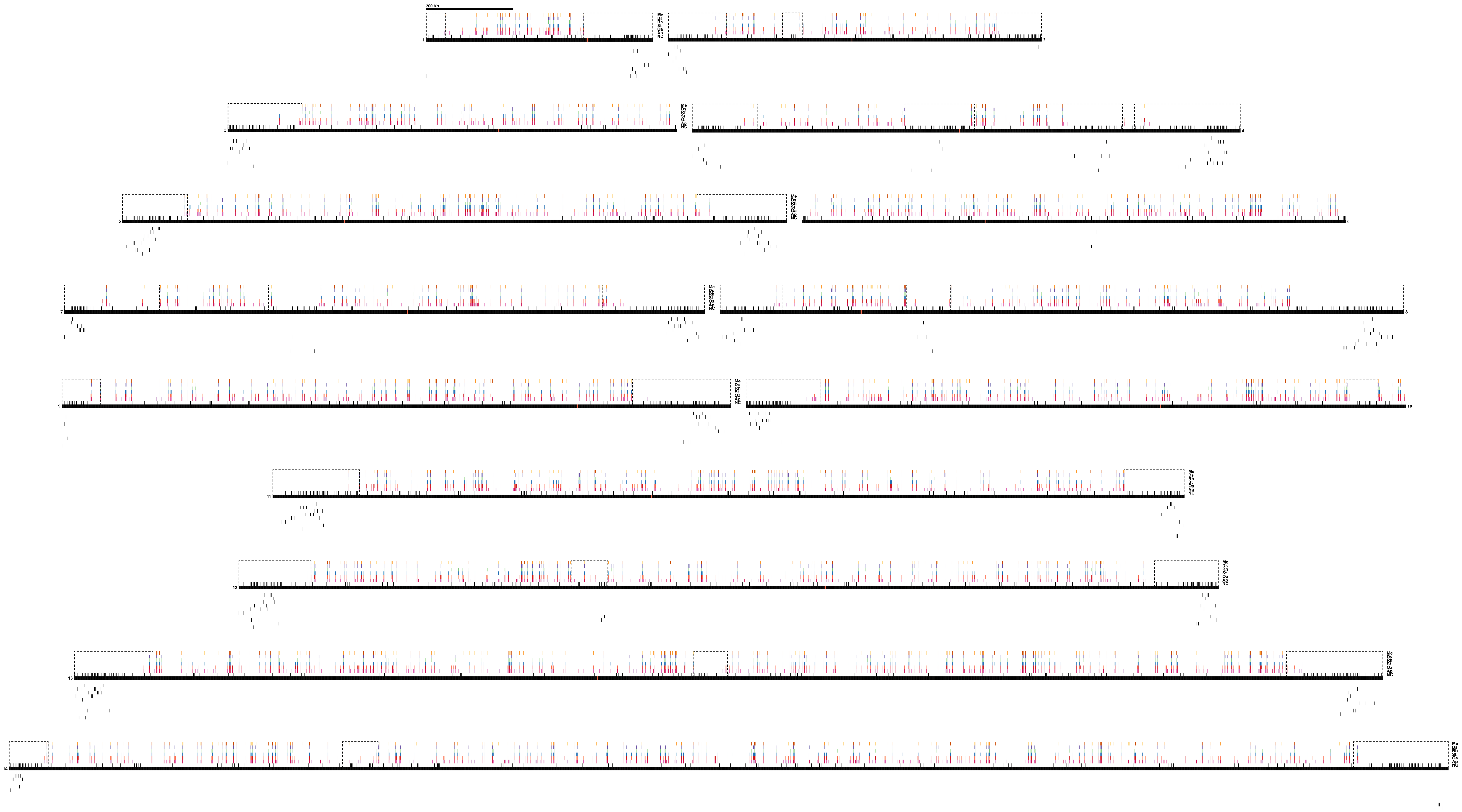
200 Kb
Me
Ds
Rh
St
Oa
Ap
NC
1
2
3
4
5
6
7
8
9
10
11
12
13
14

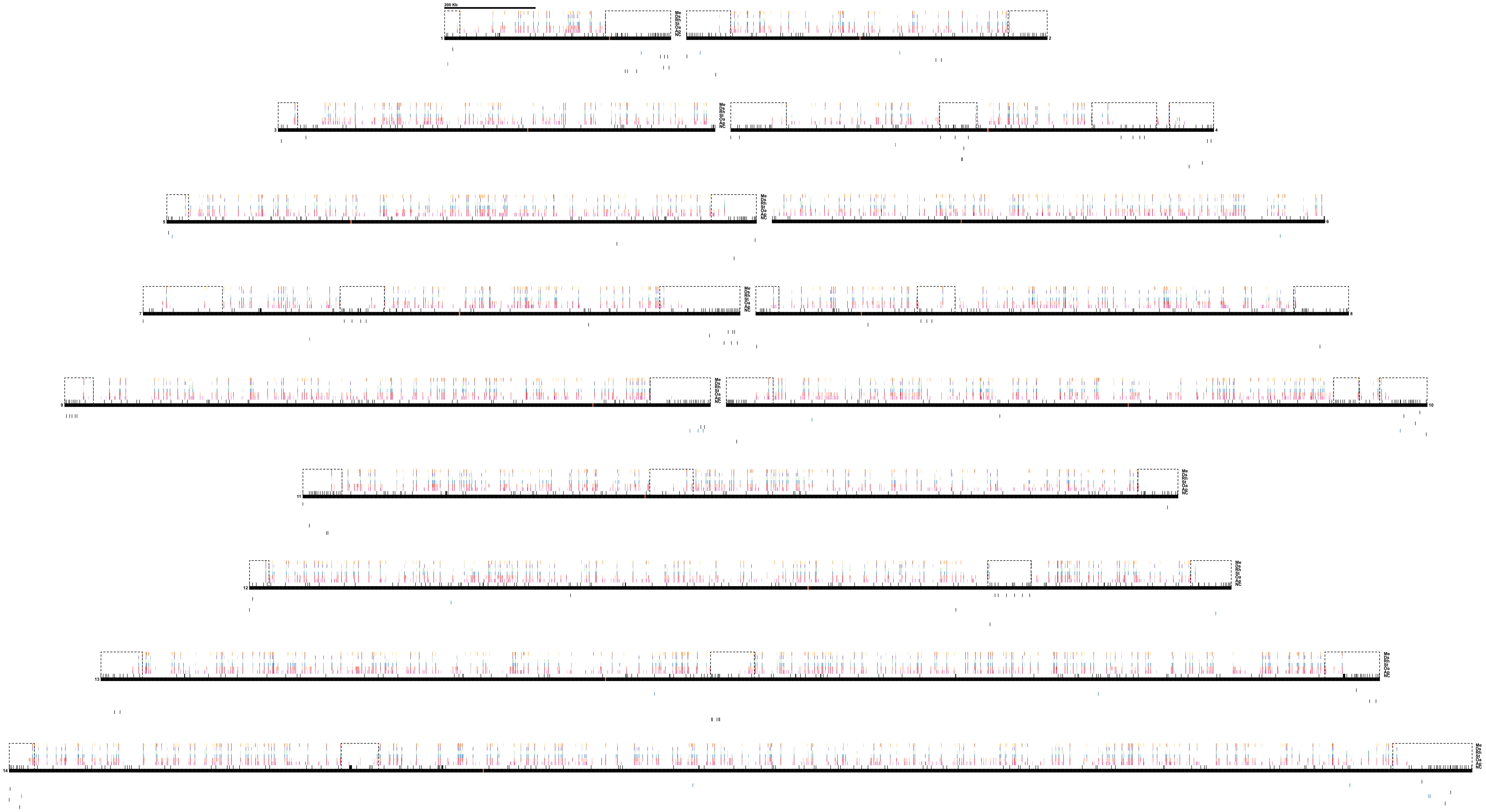

200 Kb
Me
Ds
Rh
St
Oa
Ap
NC
1
2
3
4
5
6
7
8
9
10
11
12
13
14

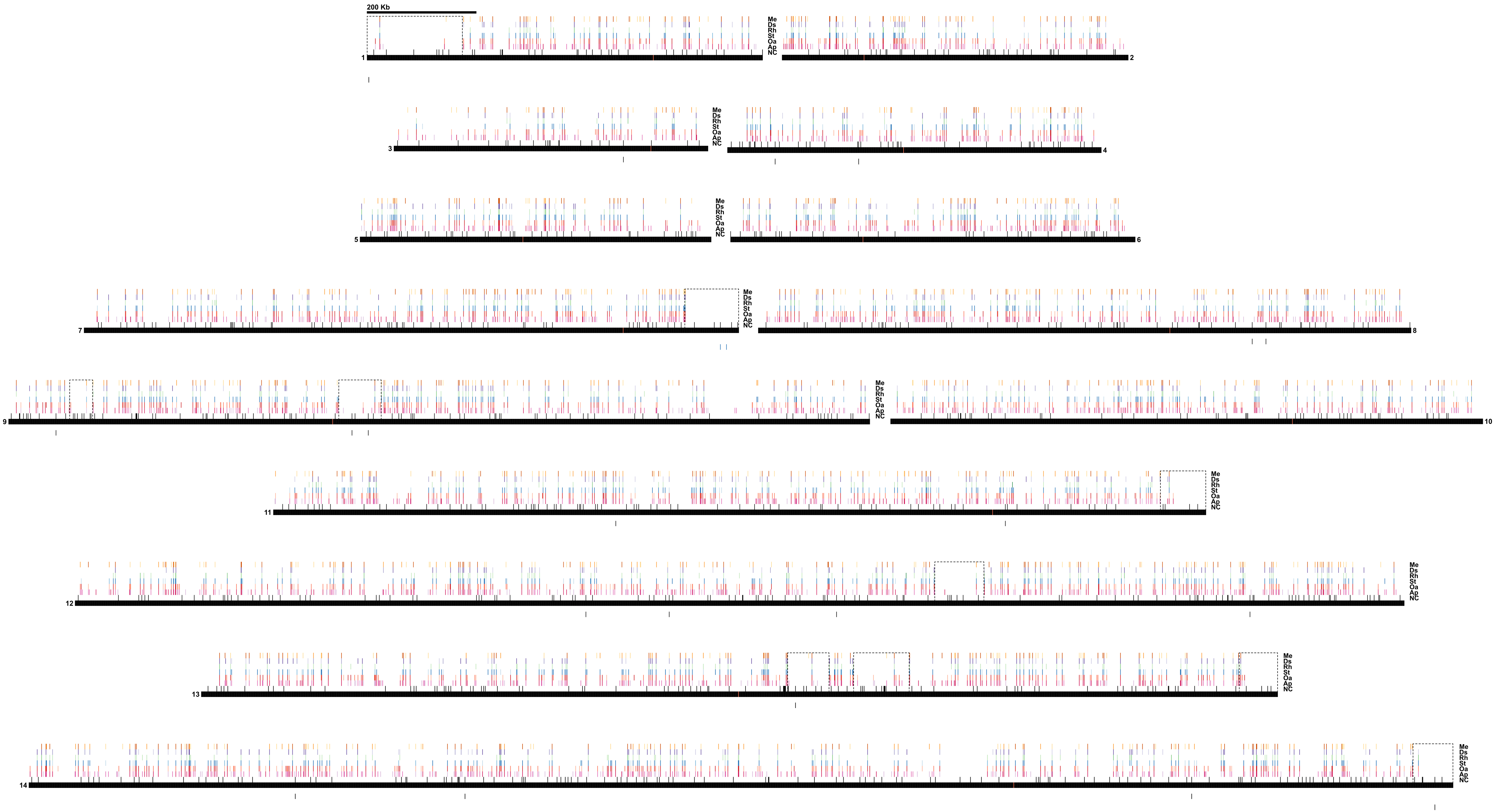
200 Kb
Me
Ds
Rh
St
Oa
Ap
NC
1
2
3
4
5
6
7
8
9
10
11
12
13
14

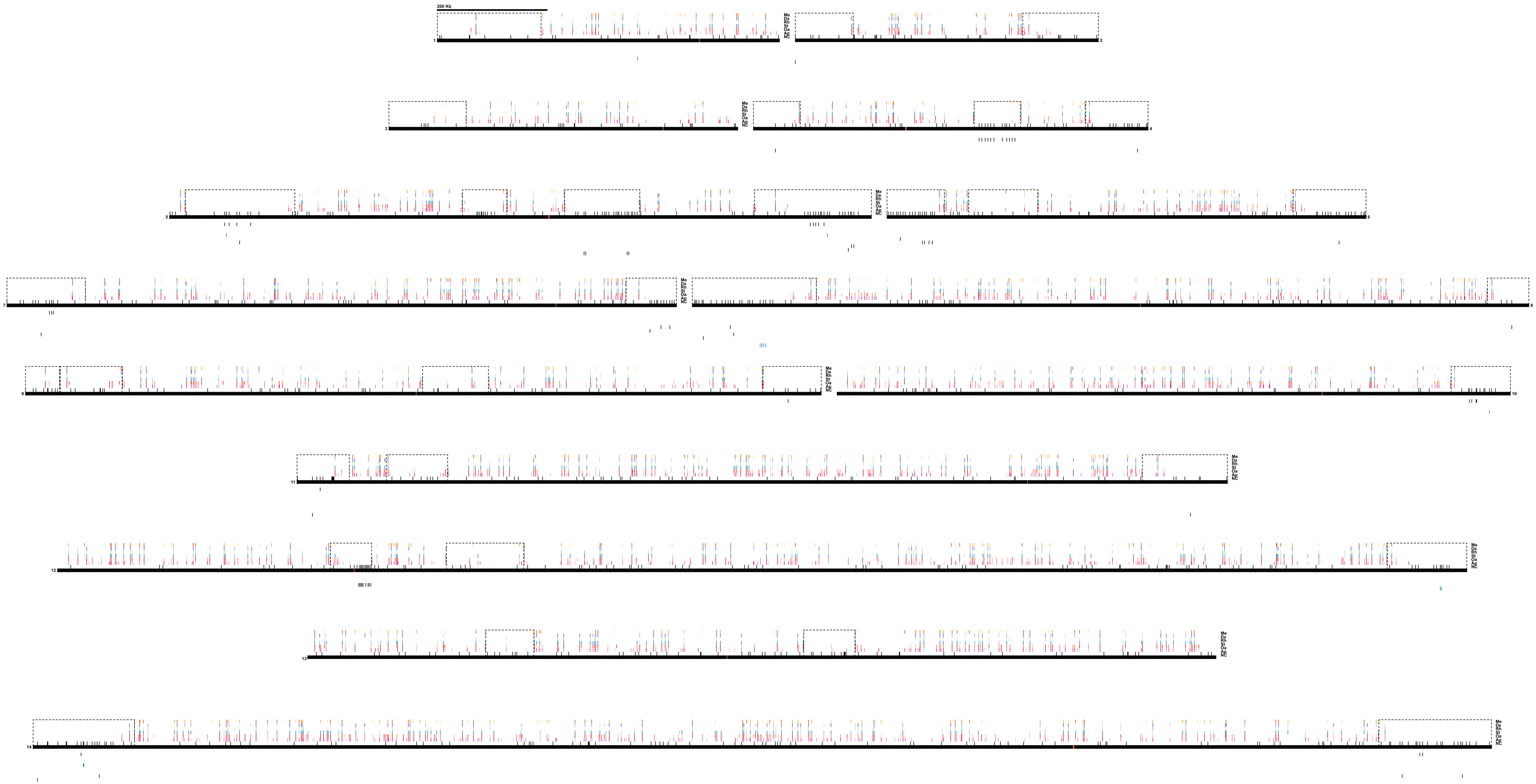
200 Kb
Me
Ds
Rh
St
Oa
Ap
NC
1
2
3
4
5
6
7
8
9
10
11
12
13
14

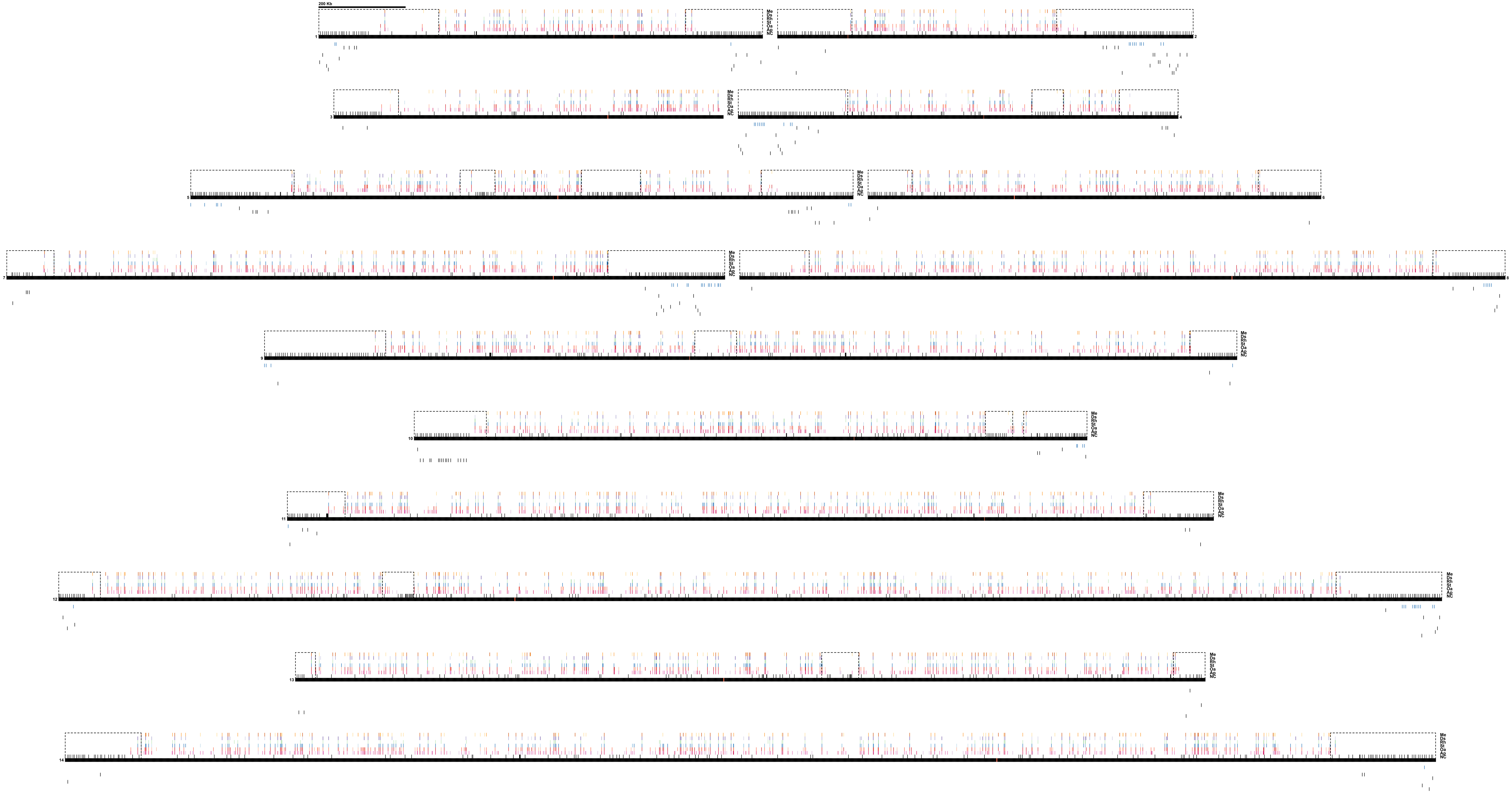
200 Kb
Me
Ds
Rh
St
Oa
Ap
NC
1
2
3
4
5
6
7
8
9
10
11
12
13
14

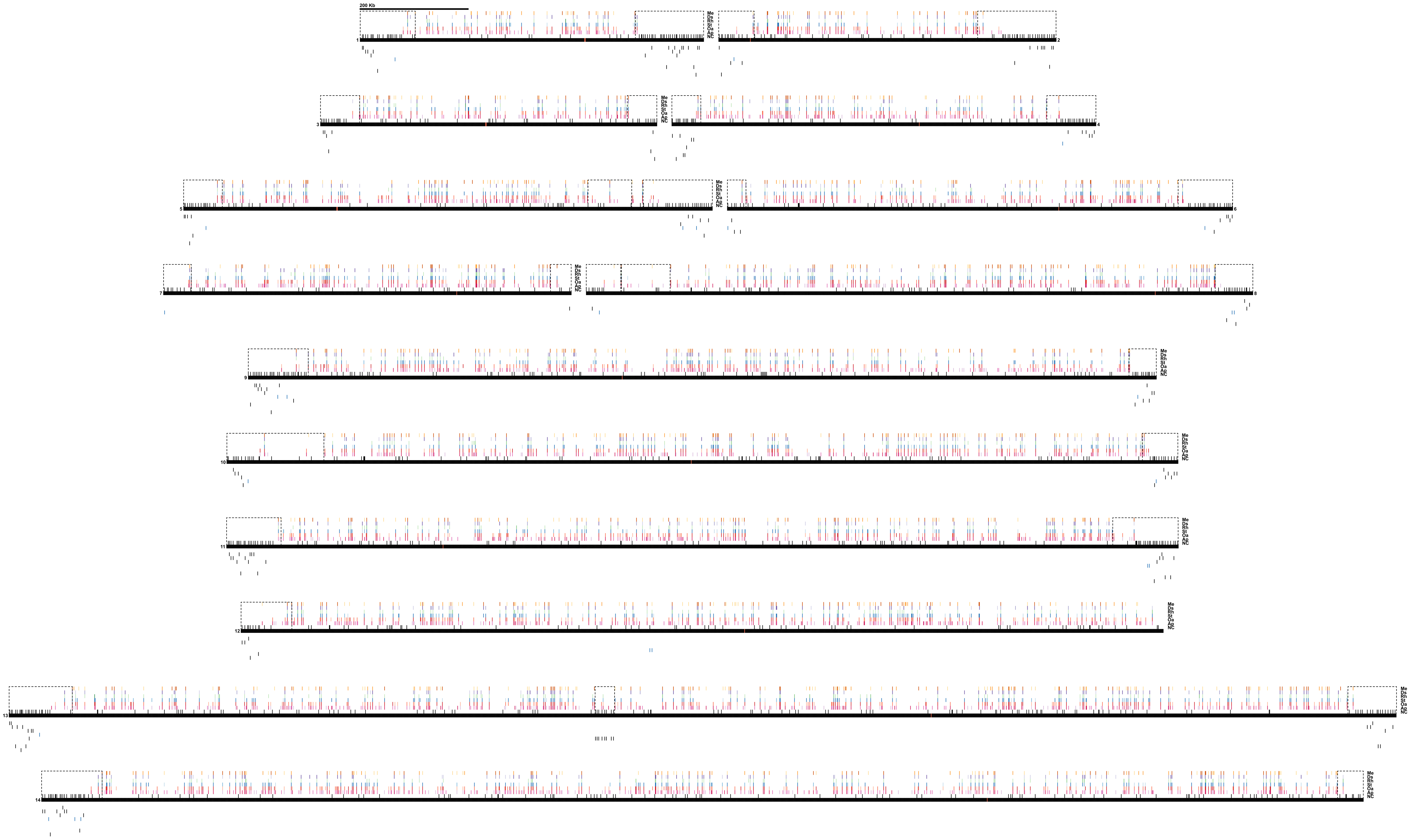
200 Kb
Me
Ds
Rh
St
Oa
Ap
NC
1
2
3
4
5
6
7
8
9
10
11
12
13
14

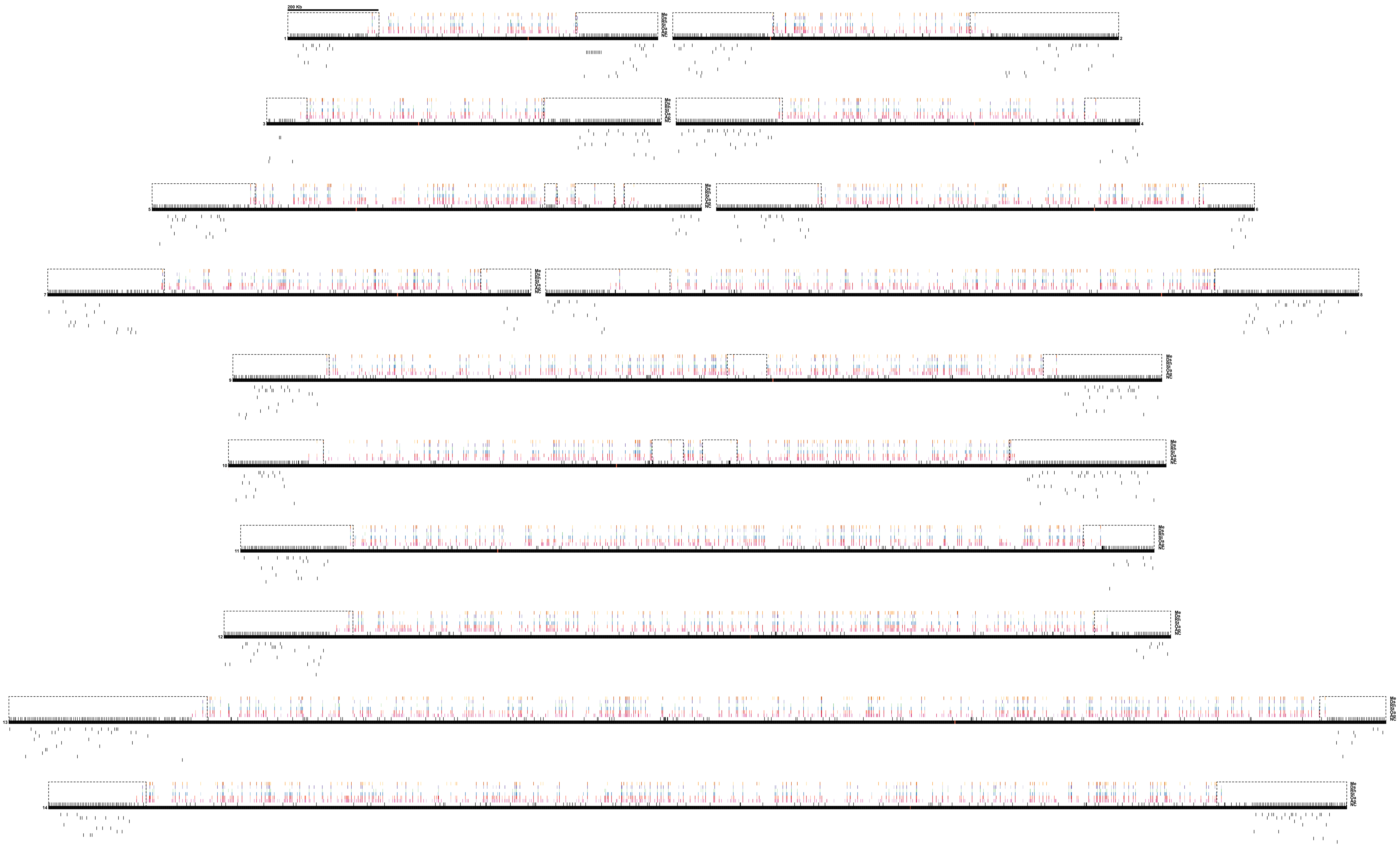
200 Kb
Me
Ds
Rh
St
Oa
Ap
NC
1
2
3
4
5
6
7
8
9
10
11
12
13
14
